# A gradual transition toward categorical representations along the visual hierarchy during working memory, but not perception

**DOI:** 10.1101/2023.05.18.541327

**Authors:** Chaipat Chunharas, Michael J. Wolff, Meike D. Hettwer, Rosanne L. Rademaker

**Affiliations:** Department of Medicine, King Chulalongkorn Memorial Hospital, Chulalongkorn University, Bangkok, Thailand; Ernst Strüngmann Institute (ESI) for Neuroscience in Cooperation with the Max Planck Society, Frankfurt, Germany; Max Planck School of Cognition, Max Planck Institute of Human Cognitive and Brain Sciences, Leipzig, Germany; Institute of Systems Neuroscience, Medical Faculty, Heinrich Heine University Düsseldorf, Germany

**Keywords:** visual perception, visual working memory, sensory recruitment, visual cortex, parietal cortex, categorization, representational similarity, representational geometry, efficient coding, RSA

## Abstract

The ability to stably maintain visual information over brief delays is central to healthy cognitive functioning, as is the ability to differentiate such internal representations from external inputs. One possible way to achieve both is via multiple concurrent mnemonic representations along the visual hierarchy that differ systematically from the representations of perceptual inputs. To test this possibility, we examine orientation representations along the visual hierarchy during perception and working memory. Human participants directly viewed, or held in mind, oriented grating patterns, and the similarity between fMRI activation patterns for different orientations was calculated throughout retinotopic cortex. During direct viewing of grating stimuli, similarity was relatively evenly distributed amongst all orientations, while during working memory the similarity was higher around oblique orientations. We modeled these differences in representational geometry based on the known distribution of orientation information in the natural world: The “veridical” model uses an efficient coding framework to capture hypothesized representations during visual perception. The “categorical” model assumes that different “psychological distances” between orientations result in orientation categorization relative to cardinal axes. During direct perception, the veridical model explained the data well. During working memory, the categorical model gradually gained explanatory power over the veridical model for increasingly anterior retinotopic regions. Thus, directly viewed images are represented veridically, but once visual information is no longer tethered to the sensory world there is a gradual progression to more categorical mnemonic formats along the visual hierarchy.

## Introduction

Holding images in mind over a brief delay is central to cognition, as it allows for the retention and manipulation of information that cannot be viewed directly. Visual working memory (VWM) recruits early visual cortex, including primary visual area V1 – as indexed by response patterns recorded with fMRI^1–8^. Being the first cortical processing site of visual inputs, the role of V1 during perception is fundamentally different from its role during visual working memory. This is because in the absence of direct visual input, mnemonic information in V1 and other early visual areas must necessarily be generated internally. Famously, sensory recruitment theory posits that higher-order frontal and parietal regions of the brain that are active throughout the working memory delay^9–20^, recruit early sensory areas in a top-down manner in order to maintain high fidelity sensory memories^21,22^. Alternatively, recurrent processes in local circuits could sustain information over a memory delay^23,24^, although such recurrenc^3,4^ y is presumably stronger in more anterior brain areas where higher pyramidal cell spine counts are believed to support increased connection strength of local circuits^25,26^. Irrespective of the exact substrate of working memory maintenance – with inputs from the external sensory world during perception, and from sources within the brain during working memory, it is unlikely that viewed and remembered visual information would be represented in an identical manner in early visual cortex^27–30^. However, it remains an open question how a cortical area like V1, specialized for processing visual inputs, actually represents visual working memories. Are working memory representations just noisier versions of perceptual representations, or do they differ in a fundamental way? Might there be representational transformations along the visual hierarchy as top-down influences play an increasingly larger role during VWM?

Recent work has claimed that early visual cortex (EVC) represents VWM information in a “sensory-like” format that is similar to representations driven by sensory inputs^1,31^, while more anterior visual areas like the Intraparietal Sulcus (IPS) represent VWM information in a format that is transformed away from the sensory driven response^31^. This claim of “sensory-likeness” in EVC comes from the fact that when participants remember an orientation, response patterns are similar to response patterns evoked by directly viewing a stimulus with the same orientation even when that viewed stimulus is not attended. In parietal cortex such cross-generalization from sensory to working memory responses fails, while memories *are* decodable when considering only the response patterns during working memory themselves (i.e., without cross-generalization). The idea that visual representations are transformed away from the “sensory-like” into a different, more abstract format during memory is further supported by work similarly using cross-generalization to decode VWM contents^32^. In this study, participants remembered one of two visually distinct features – the orientation of a grating, or the direction of a moving dot cloud. During encoding (i.e., stimulus perception), the two features evoked distinct response patterns in EVC that did not cross-generalize, likely owing to the distinct retinal inputs evoked by the two features. However, these two features share a spatial component (degrees on a circle), and during the memory delay the EVC response patterns for orientation and direction of the stimuli did successfully cross-generalize, likely owing to a “line-like” abstraction that is realizable for both visual features.

However, the possible range of to-be-remembered visual features far outstrips those that can be mapped onto a circle (and into a line). For example, surface features such as contrast or color are not readily abstracted into a “line-like” representation, let alone more complex features such as shapes or objects. Indeed, there is abundant psychophysical work showing that memory reports for numerous (surface) features are biased toward category centers^33,34,35^, indicative of categorization at the behavioral level. This means that a more general principle of abstraction remains to be uncovered. Furthermore, we know that retinotopically organized “sensory-like” representations or abstractions are not generally observed outside of early visual cortex^31,32,36,37^. An important step to investigate possible abstraction used for working memory is to consider not only *response patterns*, but also the *representational geometry*. A *response pattern* (also called a “coding scheme”^38^) simply refers to the pattern of responses that is measured during an experimental condition. Depending on the measurement technique, this could be a pattern of firing rates across multiple neurons, a pattern of BOLD responses across multiple voxels, or any other kind of response vector measured over a number of units. When it’s possible to cross-generalize from one experimental condition to another – for example from sensory to memory, or from orientation to direction – we know that the *response pattern* is similar in the two conditions. For example, the specific pattern of brain activity in response to a directly viewed stimulus with an orientation of 90° would be similar to the pattern measured when that same 90° stimulus is held in working memory. The *representational geometry* captures a lower-dimensional format of a given stimulus set, and can be invariant to changes in the underlying response patterns^39^. The pairwise distances between patterns of responses corresponding to a set of stimuli determine the geometry, meaning that even if the underlying response patterns change (e.g., they are inverted, shifted, or undergo some other transformation), the geometry can remain stable. For example, in the case of orientation the geometry may reveal that adjacent orientations (say 90° and 91°) evoke similar underlying response patterns, and that this similarity drops at increasing distances in orientation space. Such representational geometry can be shared between direct sensory input and working memory maintenance, while a sensed stimulus of 90° may nevertheless evoke a completely different response pattern compared to a 90° stimulus held in working memory. As long as the pattern distances between different orientations are the same during a sensory and memory task, so is the geometry. Indeed, we know that despite dynamics in population response patterns over time, the representational geometry of a stimulus set can remain stable^40,41^.

Here we want to examine the representational geometry during orientation working memory throughout the visual hierarchy, and see how this compares to bottom-up stimulus driven activity elicited by directly viewed stimuli. To ensure active perception of directly viewed sensory stimuli, while avoiding overlap in top-down attentional state, participants attended an orthogonal stimulus feature (contrast) during the sensory task, while holding a precise orientation in mind during the memory task. By looking at the representational geometry we can investigate the representational formats of both perception and working memory in a way that does not depend on response patterns generalizing from one condition to another. Moreover, it allows us to investigate potential systematic differences in the representational geometry between perception and working memory, even when underlying response patterns are still similar enough for successful cross-generalization. To illustrate, what if abstraction in memory happens by compressing part of the stimulus space, making it more categorical? Such compression may warp response patterns for a subset of orientations, without necessarily transforming them to a point where the coding scheme breaks down. In such a case, cross-generalization from perception to working memory could coexist with a change in the representational geometry. Thus, examining the representational geometry allows us more freedom to see if perception and working memory are truly represented similarly, or if the representational formats perhaps differ in fundamental ways.

We introduce a principled approach to investigating categorization across the visual hierarchy. First, we use representational similarity analysis (RSA) to show a clear differentiation between the geometry of perception and working memory representations for orientation in human visual cortex. Second, we model the extent to which the representational geometry is true to a simulated sensory response (the “veridical” model), or abstracted away from the sensory input (the “categorical” model). Our two models are constrained by a single principle, namely, the distribution of orientation information in the natural world^42,43^. By applying these two models to the data, we show that sensory inputs are represented in a largely veridical manner in EVC, adapted to the statistics of the natural visual world (i.e., efficient coding). By contrast, working memories are represented more categorically, in a manner predicted from the same landscape of visual input statistics, but using a higher-order metric based on how different any set of orientations may appear to the observer (i.e., their “psychological distance”). Critically, we show that during working memory the representational format becomes increasingly more categorical along the visual hierarchy, uncovering a gradient of abstraction.

## Results

To examine neural representations during perception and working memory, we analyze existing fMRI recordings from six participants who were either directly viewing oriented grating stimuli during a sensory task, or remembering orientations for later recall during a working memory task (Figure 1A). During the sensory task, participants saw each oriented grating for 9s at a time, and they had to actively detect a small reduction in contrast (500ms) that happened probabilistically (twice per 9s, meaning it could occur 0 to >2 times per trial). To avoid adaptation, grating contrast was phase reversed every 500ms. During the working memory task, participants briefly (500ms) saw a grating and remembered its orientation for 13s. On two-thirds of memory trials a visual distractor (filtered noise, or another grating with uncorrelated orientation) was presented during the middle portion of the delay (for 11 seconds). Because there was little-to-no quantifiable difference between the different trial types (see ref^31^), we analyzed data from all delays combined. For all analyses we use average voxel responses from 4.8–9.6s and 5.6–13.6s after stimulus onset for the sensory and memory task, respectively (replicating the time-windows used in the original publication^31^). For the working memory task in particular, this choice of time-window is based on the observation that (1) the BOLD response evoked by the to-be-remembered stimulus is back to baseline ∼6s post stimulus onset in these data^31^, (2) decoding is possible for every single TR during the delay^31^ well beyond the stimulus-evoked BOLD response and despite poorer signal-to-noise of single TR data, and (3) much prior work has shown that stimulus-evoked BOLD alone is insufficient for stimulus information to persist into the memory delay^1,2,4,8,44,45^.

**Figure 1:**
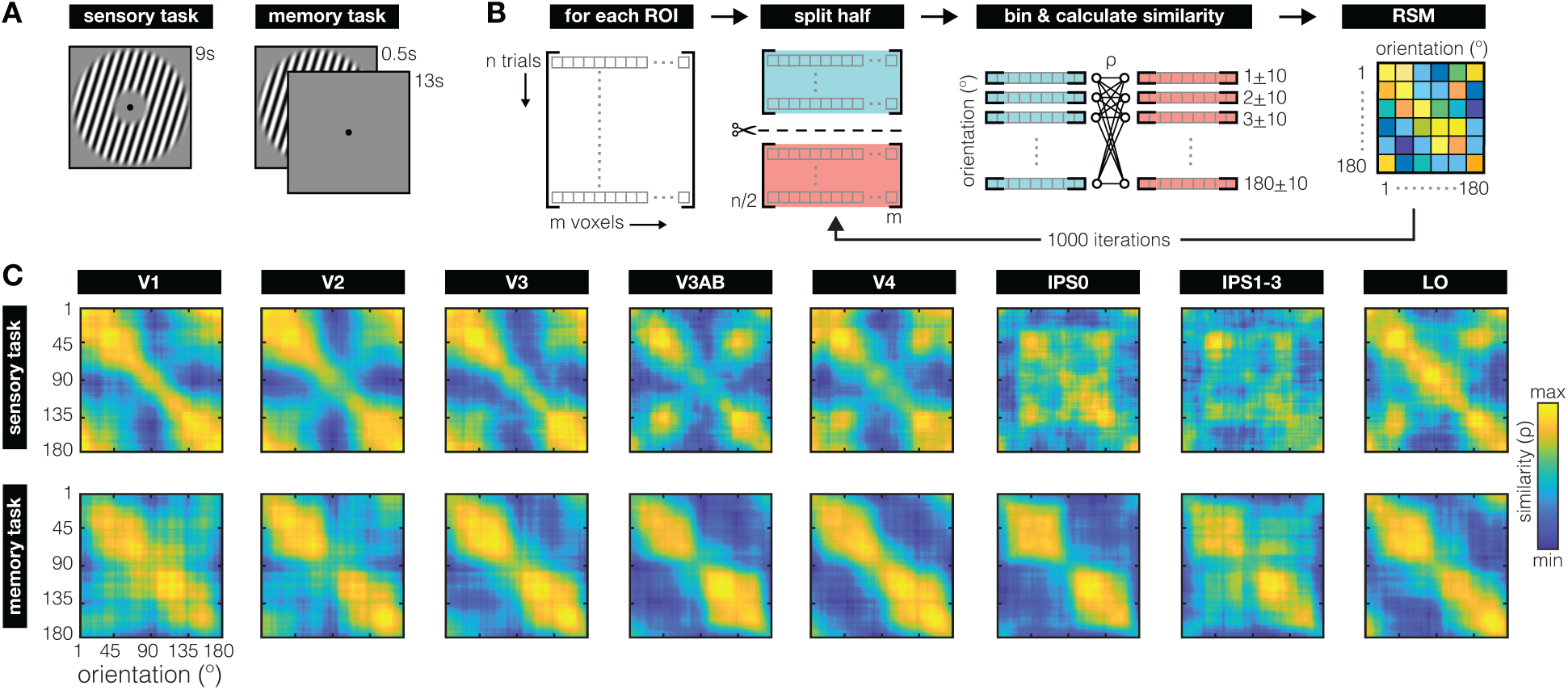
Task and main analysis. **(A)** For the sensory task (left), participants viewed a randomly oriented grating for 9 seconds per trial (contrast phase-reversing at 5 Hz) and reported instances of contrast dimming. For the working memory task (right), participants remembered a briefly presented (500 ms) randomly orientated grating for 13 seconds, until a 3 second recall epoch (not depicted). **(B)** For each Region of Interest (ROI) we employed a split-half randomization procedure to create a Representational Similarity Matrix (RSM) for each participant. On each randomization fold, voxel patterns from all trials (300–340 for sensory, 324 for memory) were randomly split in half. For each half of trials, we averaged the voxel patterns for every degree in orientation space within a + 10° window. This resulted in 180 vectors with a length equal to the number of voxels for each split of the data. We then calculated the similarity between each vector (or degree) in one half of the data, to all vectors (or degrees) in the second half of the data, using a Spearman correlation coefficient. This resulted in a 180x180 similarity matrix on each fold. This randomization procedure was repeated 1000 times to generate the final RSM for each ROI and each participant. Across all folds, RSM’s are near-symmetrical around the diagonal, give-or-take some cross-validation noise. **(C)** Representational geometry of orientation during the sensory (top row) and working memory (bottom row) tasks, for retinotopically defined ROI’s (columns) across all participants. During the sensory task, the clear diagonal pattern in early visual areas V1–V3 indicates that orientations adjacent in orientation space are represented more similarly than orientations further away. During the memory task, similarity clusters strongly around oblique orientations (45° and 135°), contrasting starkly with the similarity patterns during perception. Note that the diagonal represents an inherent noise-ceiling, due to the cross-validation procedure used. This noise ceiling shows inhomogeneities across orientation space, demonstrating how certain orientations may be encoded with more noise than others. RSM’s are scaled to the range of correlations within each subplot to ease visual comparison of representational structure between sensory and memory tasks for all ROI’s (exact ranges are shown in Supplementary Figure 3). For early ROI’s (V1–V4), only visually responsive voxels are included in the analysis. Throughout, 0° (and 180°) denotes vertical, and 90° denotes horizontal.

We use cross-validated representational similarity analysis (RSA) – a method that projects neural activation patterns into an abstract space that describes the stimulus (here: orientation)^41, 46–48^. Specifically, we created, for each visual cortical Region of Interest (ROI) and each of the two tasks (sensory and memory), a matrix capturing the similarity between the neural response patterns to all possible orientations using cross-validated correlations (Figure 1B). To illustrate: When two orientations are represented in a very similar manner, the pattern of voxel responses to the first orientation will correlate strongly with the pattern evoked by the second orientation. Conversely, for orientations with very distinct representations, correlations will be low. Because orientation space is continuous, physically similar orientations (e.g., 10° and 11°) will likely correlate more strongly than physically dissimilar orientations (e.g., 10° and 40°) in areas of the brain that encode orientation information.

The representational similarity matrices (RSM’s) constructed for our visual ROI’s (Figure 1C; Supplementary Figure 1) show striking qualitative differences between how orientation is represented during perception (in the sensory task), and working memory (in the memory task). During the sensory task, early visual areas V1–V3 show a strong diagonal component and relatively higher degree of similarity around cardinal orientations (180° in particular, which is vertical), with notable transformations away from this representational geometry primarily along the dorsal stream (V3AB and IPS). During the memory task, there is a prominent clustering of representational similarity around oblique orientations (45° and 135°). This clustering seems to increase along the visual hierarchy and appears most pronounced in area IPS0. These results do not depend on the specific way we bin orientations in our RSA (Supplementary Figure 2).

To quantify the results in Figure 1C, we contrast two possible models – the “veridical” and the “categorical” model. The veridical model intends to capture how orientations are represented in a manner that faithfully reflects early visual processing of signals from the external world. The categorical model uses a higher-level concept – the “psychological distance” between orientations – as a basis for abstracting a physically continuous space into discrete categories. Importantly, these two models are jointly constrained by the known distribution of orientation information in the natural world, which has a higher occurrence of cardinal compared to oblique orientations (Figure 2A; ref^42^). According to the “efficient coding” hypothesis, this inhomogeneity in orientation input statistics leads to adaptive changes in the sensory system, with relatively more neural resources dedicated to cardinal compared to oblique orientations^49–53^. As a result, observers demonstrate a higher resolution (e.g., improved discriminability) around cardinal orientations – a phenomenon paradoxically known as the “oblique effect”^54–56^. Orientation reports also tend to be biased away from cardinal axes^43,57,58^. Together, this suggests a distinct role for cardinal orientations, both with respect to precision and bias.

**Figure 2:**
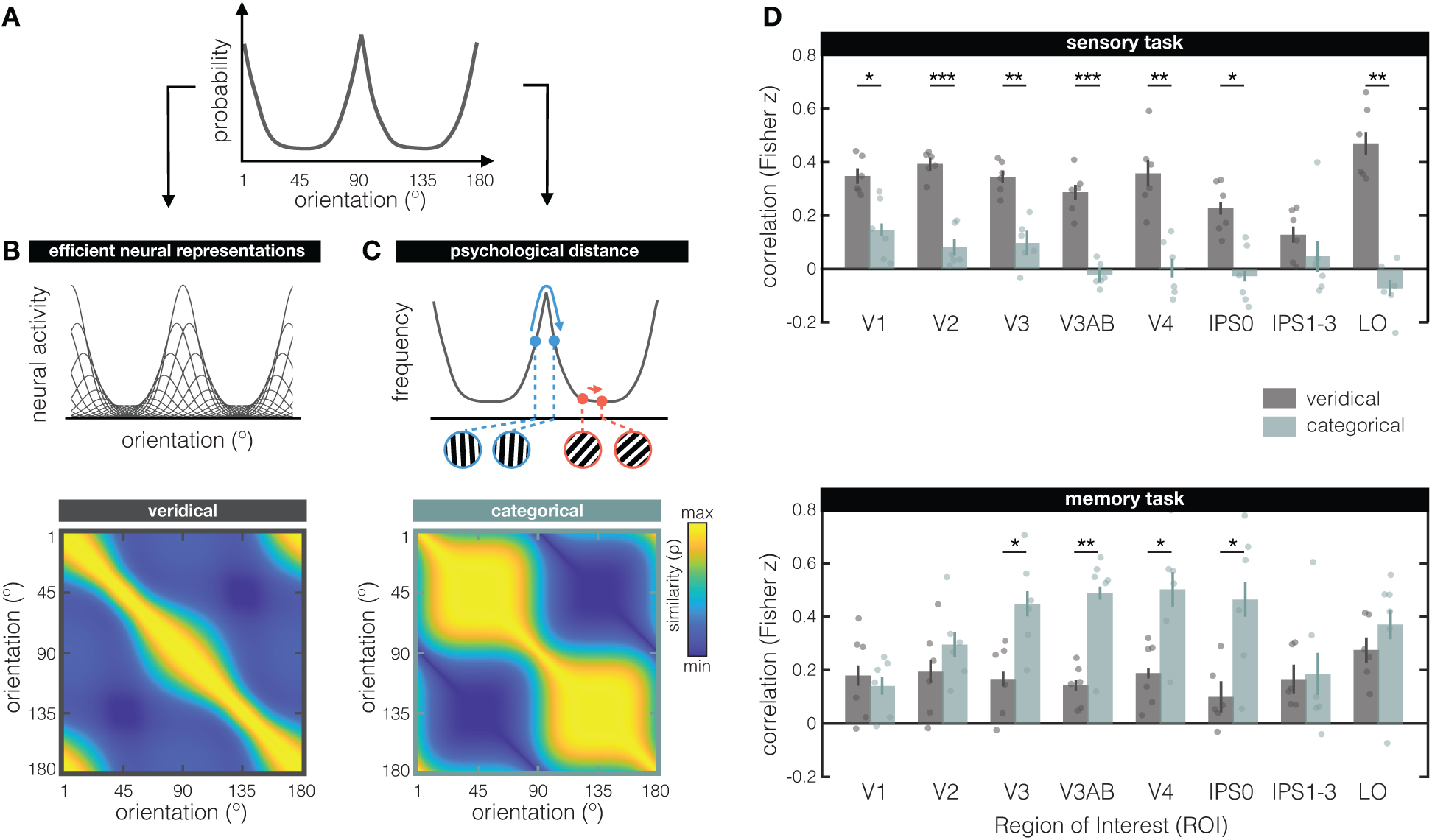
Modeling the representational similarity of perceived and remembered orientations. **(A)** The distribution of visual orientation in the natural world is inhomogeneous, with higher prevalence of orientations closer to cardinal (90° & 180°) compared to oblique (45° & 135°). The function shown here approximates these input statistics, and is used to constrain both the veridical (in **B**) and categorical (in **C**) models. **(B)** The veridical model is based on the principle of efficient coding – the idea that neural resources are adapted to the statistics of the environment. We model this via 180 idealized orientation tuning functions with amplitudes scaled by the theoretical input statistics function (the top panel shows a subset of tuning functions for illustrational purposes). A vector of neural responses is simulated by computing the activity of all 180 orientation-tuned neurons to a given stimulus orientation. Representational similarity is calculated by correlating simulated neural responses to all possible orientations, resulting in the veridical model RSM (bottom panel). Note that while we chose to modulate tuning curve amplitude, there are multiple ways to warp the stimulus space (e.g., by applying non-uniform changes to gain, tuning width, tuning preference, etc.^39,43^). **(C)** In the categorical model, categorization is based on people’s subjective experience of relative similarity between orientations in different parts of orientation space: If orientations in part of the space appear quite similar, they are lumped together into the same category, while the most distinctive looking orientations serve as category boundaries. This is quantified via the “psychological distance” – the sum of derivatives along the input statistics function between any pair of orientations (see top panel). The insert shows an example of orientation-pairs near cardinal (in blue) and oblique (in red) that have the same physical distance, but different psychological distances. The psychological distance between each possible pair of orientations yields the categorical model’s RSM (bottom panel). **(D)** Fits of the veridical (grey) and categorical (teal) models for the sensory (top) and memory (bottom) tasks. During the sensory task, the veridical model better explains the data compared to the categorical model in almost all visual ROI’s (except IPS1–3), indicating a representational scheme that is largely in line with modeled early sensory responses. During the memory task, the categorical model gains increasingly more explanatory power over the veridical model along the visual hierarchy, and explains the data significantly better in V3, V3AB, V4, and IPS0. The Fisher transformed semi-partial correlations (on the y-axis) represent the unique contribution of each model after removing the variance explained by the other model via semi-partial correlations. Dots represent individual participants, and errorbars represent + 1 within-participant SEM. Asterisks indicate the significance level of post-hoc two-sided paired-sample t-tests (*p < 0.05; **p < 0.01; ***p < 0.001) comparing the two models in each ROI.

The idea behind the veridical model is that region-wide orientation representations emerge from low-level neural responses to sensory inputs. Specifically, the starting point for this model is a set of idealized tuning functions that tile orientation-space (Figure 2B, top). The amplitude of each orientation tuning function is scaled by the estimated frequency of occurrence for that orientation in the natural world (i.e., scaled by the theoretical “input statistics” function, see Figure 2A and Methods). Therefore, tuning functions closer to cardinal orientations have relatively higher amplitudes than those closer to obliques. We modulate tuning curve amplitude (and not other properties like tuning width or density) because of known amplitude differences in the fMRI signal for cardinals compared to obliques^59^. For any given stimulus orientation, we can simulate a vector of neural responses by reading out the hypothesized activity from every idealized neural tuning function. Such a simulated response vector can be correlated against simulated responses to all other possible stimulus orientations (analogous to the approach in Figure 1B), to arrive at the veridical model RSM (Figure 2B, bottom). Specifically, to preserve the BOLD amplitude differences mentioned above, we use Pearson correlations to generate the model RSM. Thus, here we model a veridical early visual representation by accounting for known inhomogeneities of orientation space, based on the principle of efficient coding^43^. Note that our veridical model is a direct consequence of the choice to use amplitude modulation (instead of e.g., tuning width) as well as correlation (instead of e.g., Euclidian distance), and that there are multiple other possible ways to implement inhomogeneities across orientation space and the idea of efficient coding^39,43,60^.

The idea behind the categorical model is to discretize the physically continuous orientation space into plausible higher-level psychological categories. A principled way to categorize orientation is to again rely on the input statistics function (Figure 2A), and consider how inhomogeneities in orientation space might affect more experiential measures such as perceptual similarity^61^. For example, in parts of the orientation space with high resolution (near the cardinals), two physically similar orientations (e.g., 4° apart) can look clearly different from one another, while orientations with the same physical similarity in parts of orientation space with lower resolution (near the obliques) can look indistinguishable. We formalize this “psychological distance” as the distance between any pair of orientations along the theoretical input statistics function (Figure 2C, top). Returning to our example, two near-cardinal orientations (e.g., 88° and 92°) will have a larger psychological distance (i.e., are relatively far apart along this function) compared to two near- oblique orientations (e.g., 43° and 47°) (compare the Figure 2C grating inserts in blue versus red, respectively). The psychological distances between all possible orientations make up the categorical model RSM (Figure 2C, bottom). This RSM shows how orientations in one category (bound by two cardinals) are represented similarly to one another, but dissimilarly from orientations in a second category (on the other side of the cardinals). Thus, here we model how orientation representations can be categorized based on where an orientation is relative to cardinal – the cardinal axes effectively serving as category boundaries.

How well can the representational geometry during the sensory and memory tasks (Figure 1C) be explained by our veridical (Figure 2B) and categorical (Figure 2C) models? Because our two models are not independent, we evaluated the correlation of each model to the data after first removing the variance explained by the other model, which is also known as a semi-partial correlation. Specifically, to look at the unique contribution of the veridical model in explaining the data, we first remove the variance of the categorical model from the veridical model, and then correlate the residuals to the data RSM’s (and vice versa for the categorical model; see Methods). We apply a Fisher transformation to the resulting correlation values to normalize their distribution, and better allow for statistical testing. Transformed correlations are shown in Figure 2D for our sensory (top) and memory (bottom) tasks. Importantly, these results do not depend on the specific fitting approach, and replicate when we fit each model directly to the data (without first taking the residuals), and also when we use a general linear model to simultaneously fit both models to estimate their beta weights (Supplementary Figure 4).

During the sensory task, the veridical model does an overall better job at explaining representational similarity than the categorical model (main effect of model: F_(1,5)_ = 76.05, p = 0.0003). This advantage is not the same in all ROI’s (interaction of model x ROI: F_(7,35)_ = 4.158, p = 0.002), with post-hoc tests showing a difference between the two models in all ROI (except IPS1–3). The fact that the veridical model outperforms the categorical model during perception in the sensory task helps validate our modeling approach, given that the veridical model is founded on what we know about early bottom-up sensory processing.

During the memory task, the categorical model better explains representational similarity than the veridical model (main effect of model: F_(1,5)_ = 11.4, p = 0.0198). The extent to which the categorical model outperforms the veridical model differs across ROI’s (interaction of model x ROI: F_(7,35)_ = 3.048, p = 0.013), with the categorical model explaining increasingly more of the representational geometry along the visual hierarchy. A significant difference between the two models emerges in area V3, and persists until area IPS0. These results corroborate the qualitative categorization already apparent from the memory task RSM’s in Figure 1C, and reveal a posterior-to-anterior gradient in terms of categorization strength during working memory, but not perception (interaction of ROI x task x model: F_(7,35)_ = 3.226, p = 0.0095). Finally, we also confirmed that the presence of distractors during the delay did not impact the pattern of results in the memory task (Supplementary Figure 5).

To verify that our modeling results do not critically depend on the exact shape of the theoretical input statistics function in Figure 2A, we next used behavioral error from an independent psychophysical experiment to constrain both our models (Figure 3A). A new set of 17 participants each completed 1620 trials of an orientation recall task. For every possible stimulus orientation that was shown (1°–180° in steps of 1°), we calculate the absolute mean recall error across all participants. With 27540 total trials in the experiment, and a sliding window of 3° for more reliable estimates, absolute errors are based on 459 trials for every possible stimulus orientation. We chose the mean absolute error as it takes both response variance and response bias into account. As expected, recall error is inhomogeneous along orientation space, with pronounced differences between cardinals and obliques (Figure 3A). Specifically, recall is more accurate around cardinal compared to oblique orientations. We generated the veridical and categorical models anew from this psychophysical input function (Figure 3B), and again fit both models to our empirical RSM’s to see how well they explained the data (Figure 3C). We replicated the difference between the sensory and memory tasks, and how their representational geometries are better explained by the veridical and categorical models, respectively (interaction ROI x task x model: F_(7,35)_ = 4.712; p = 0.001). During the sensory task, the veridical model outperformed the categorical model (main effect of model: F_(1,5)_ = 52.55, p = 0.001), and this advantage differed significantly between ROI’s (interaction of model x ROI: F_(7,35)_ = 2.984, p = 0.015). During the memory task, the categorical model outperformed the veridical model (main effect of model: F_(1,5)_ = 10.6, p = 0.023), and explained increasingly more of the representational geometry in some ROI’s than in others (interaction of model x ROI: F_(7,35)_ = 4.539, p = 0.001).

**Figure 3:**
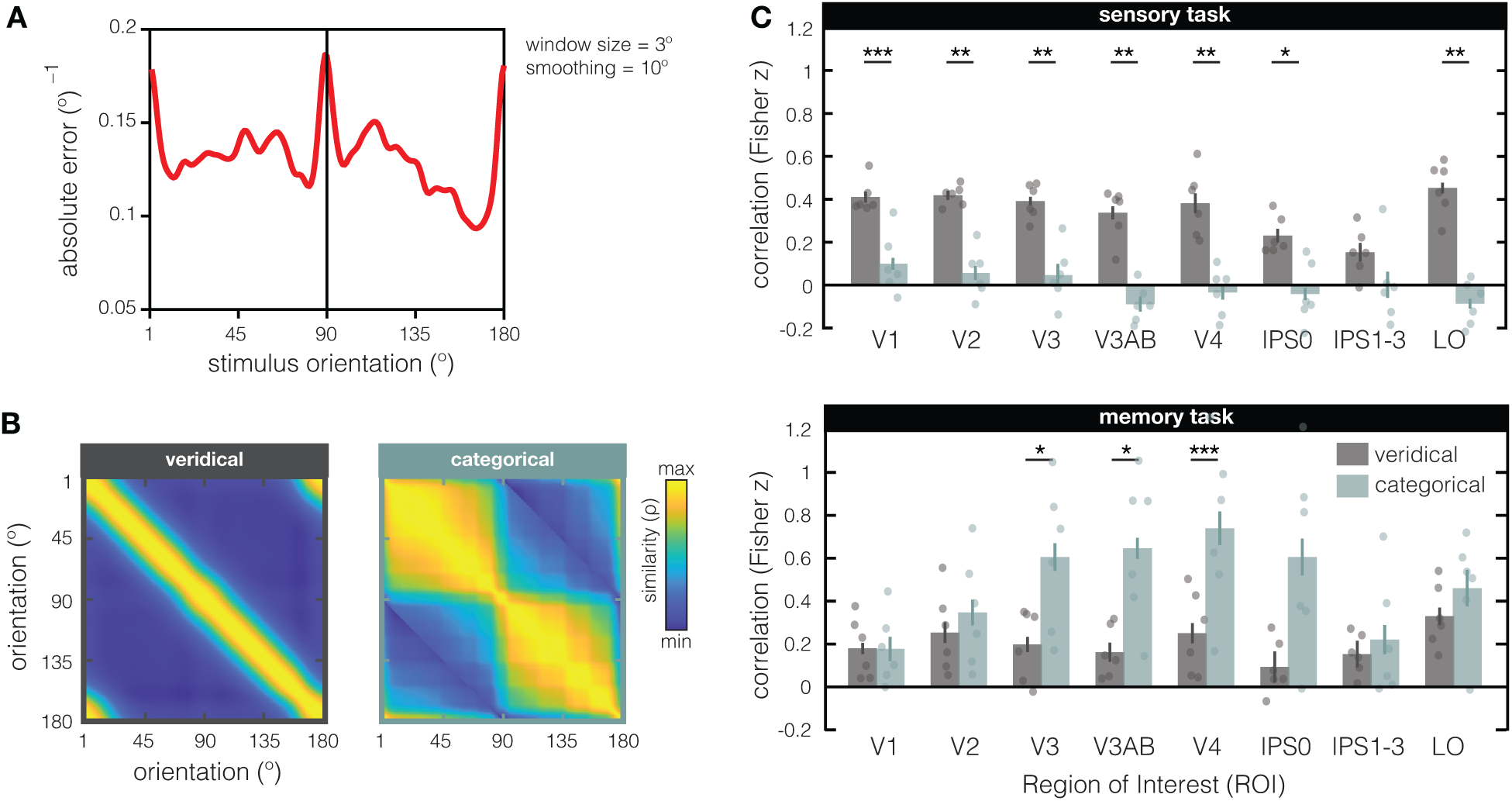
Generating and fitting the veridical and categorical models based on independent behavioral data. **(A)** During an independent psychophysical examination, a new set of participants (N=17) reported the orientation of briefly presented (200ms) and remembered (2s) single gratings by rotating a response dial with a computer mouse (i.e., via method-of-adjustment). For each possible stimulus orientation in the experiment (+1°), we calculated the mean absolute response error across all participants, and smooth the resulting function (Gaussian, over 10°). The absolute error^−1^ (y-axis) is plotted against the stimulus orientation shown to participants. From this psychophysical input function, the veridical and categorical models were generated as previously described (see Figure 2B & 2C). **(B)** Veridical and categorical models generated from the psychophysical input function (in **A**). **(C)** Fits of the veridical (in grey) and categorical (in teal) models based on the independent psychophysical data. During the sensory task (top), the veridical model better explains the data compared to the categorical model in all visual ROI’s except for IPS1–3. During the memory task (bottom), the categorical model better explains the data compared to the veridical model in V2, V2AB, and V4 (and marginally better in IPS0 with p = 0.053). The Fisher transformed semi-partial correlations (on the y-axis) represent the unique contribution of each model after removing the variance explained by the other model. Dots represent individual participants, and errorbars represent + 1 within-participant SEM. Asterisks indicate the significance level of post-hoc two-sided paired-sample t-tests (*p < 0.05; **p < 0.01; ***p < 0.001) comparing the two models in each ROI.

How might we reconcile the observed *differences* in representational geometry between the sensory and memory tasks, with the *overlap* in coding schemes that is evident from the ability to cross-generalize from the sensory to the memory task in EVC^31^? To directly relate these two analysis approaches, we modified the typical RSA approach by correlating response patterns from every *perceived* orientation in the sensory task to the response patterns from every *remembered* orientation in the memory task, in what we call “across-task RSA” (Figure 4A). As expected, our across-task RSM of V1 shows a clear diagonal component that is indicative of overlap in response patterns between perception and working memory, and the ability to cross-generalize. Cross-generalization does not work in IPS, replicating what we know from previous analysis of these data^31^. Importantly, this approach also shows how the coding scheme for orientation during the memory task is warped with respect to coding scheme during the sensory task – response patterns for orientations held in working memory are biased towards what would be patterns associated with obliques during perception (Figure 4B). We validate our “across-task RSA” approach against a conventional multivariate analysis approach known as the inverted encoding model (or “IEM”^62^). First, we take into account the predicted gradual drop in representational similarity for orientations at increasing distances from the remembered orientation by creating a “correlation profile” (the sum of correlations between patterns for the remembered and perceived orientations, as shown in grey on top of the panels in Figure 4B, and for all ROI’s in Figure 4C). By collapsing the data based on “same” versus “increasingly different” orientations, we’re in essence performing decoding, but by using correlation. A more “peaked” correlation profile indicates more information about the remembered orientation, which we quantify using a previously established fidelity metric (as in ^31^, see Methods). Finally, we show that this “RSA fidelity” aligns closely with the same fidelity metric applied to results from an IEM (Figure 4D).

**Figure 4:**
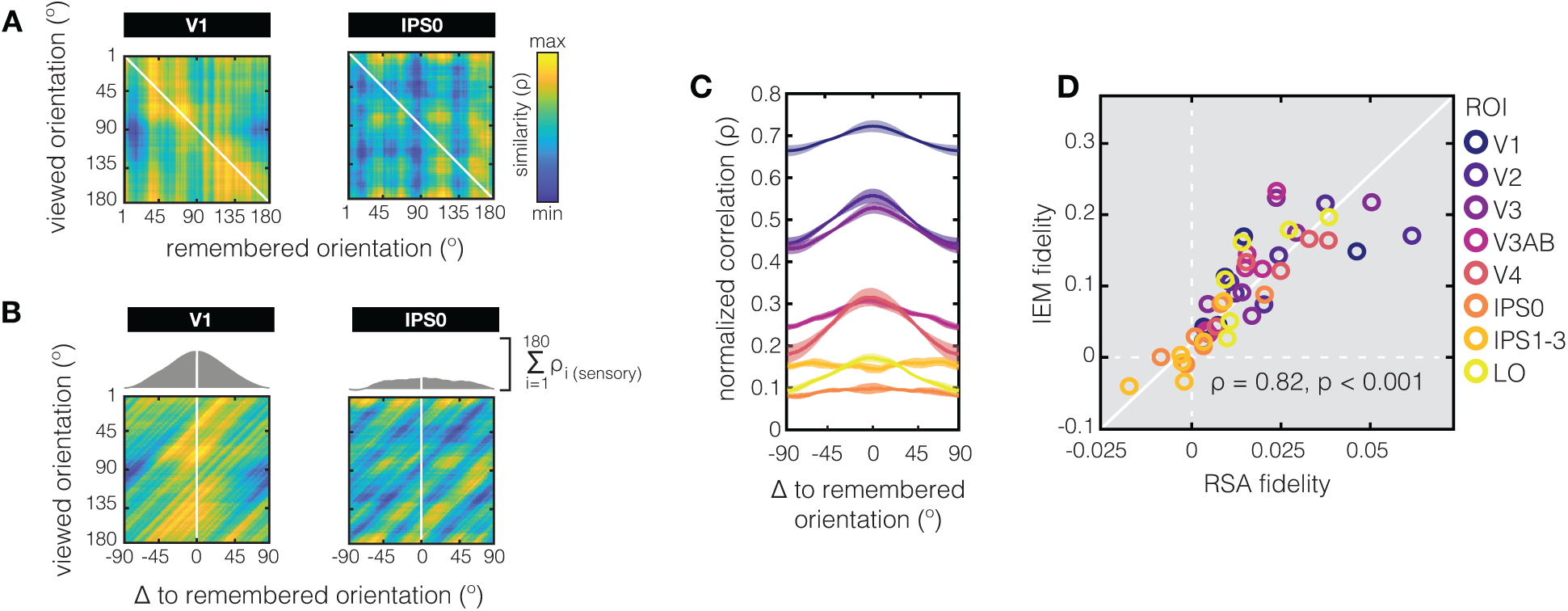
Ability to cross-decode using RSA. **(A)** Using across-task representational similarity analysis, we directly compare orientation response patterns recorded during the sensory task (y-axis), to those measured during the memory task (x-axis). Here we show V1 (left subplot) and IPS0 (right subplot) as example ROI’s. The across-task RSM in V1 shows a clear diagonal component, indicating similar response patterns for specific orientations in the sensory and memory tasks. In IPS0 such pattern similarity for matching orientations in the sensory and memory tasks is less evident. **(B)** We want to quantify the extent to which orientations held in working memory evoke response patterns that overlap with response patterns from those same orientations when viewed directly, and how this similarity drops at larger distances in orientation space. First, we center our across-task RSM’s on the remembered orientation (notice the x-axis), and then take the sum of correlations relative to the remembered orientation (plotted on top of the across-task RSM’s in grey). We call this the “correlation profile” of the remembered orientation. In V1 we see that correlations are highest between response patterns from matching perceived and remembered orientations (0° on the x-axis), explaining the ability to cross-decode between sensory and memory tasks as demonstrated in previous work (e.g.,^1,31^). By contrast, IPS0 shows a much flatter correlation profile. **(C)** Correlation profiles for all retinotopic ROI’s in our study, obtained by performing across-task RSA (left panel). Most ROI’s show a peaked correlation profile, indicative of shared pattern similarity between the same orientations when directly viewed and when remembered. The different offsets along the y-axis for different ROI’s reflect the overall differences in pattern similarity in different areas of the brain, with pattern similarity being highest in area V1. Shaded areas indicate + 1 SEM **(D)** To validate the ability to cross-decode using RSA, we directly compare this new approach (x-axis) to the multivariate analysis performed by Rademaker et al. in 2019 (y-axis). The latter used an inverted encoding model (IEM) that was trained on the sensory task, and tested on the delay period of the memory task. Both the correlation profiles from RSA, and the channel response functions from IEM yield more-or-less peaked functions over orientation space (relative to the remembered orientation) that can be quantified using the same fidelity metric (i.e., convolving with a cosine). Here, we show a high degree of consistency between the fidelity metrics derived with both approaches, and successful cross-generalization from the sensory to the memory task (as indexed by >0 fidelities) in many ROI’s. Each color represents a different ROI, and for each ROI we plot each of the six participants as an individual dot.

Thus far, we examined the structure of orientation representations during perception and working memory in individual visual ROI’s, and find that orientation representations differ between sensory and memory tasks. We also find that representational geometry *within the same task* is captured by our models to varying extents in different ROI’s. To move beyond specific patterns of information in local ROI’s (e.g., Figure 1C; Supplementary Figure 1), and more formally assess representational geometries across visual cortex, we use a 2^nd^ level RSA. In this analysis, similarity between different visual cortical areas is calculated by correlating the RSM from every ROI with that of every other ROI. To this end, we use more fine-grained ROI’s than in previous analyses, allowing us to look at dorsal versus ventral streams, as well as subregions of IPS and Lateral Occipital (LO) cortex. We evaluate how orientation information is represented across visual cortex in this manner separately for the sensory task and the memory task (Figure 5).

**Figure 5:**
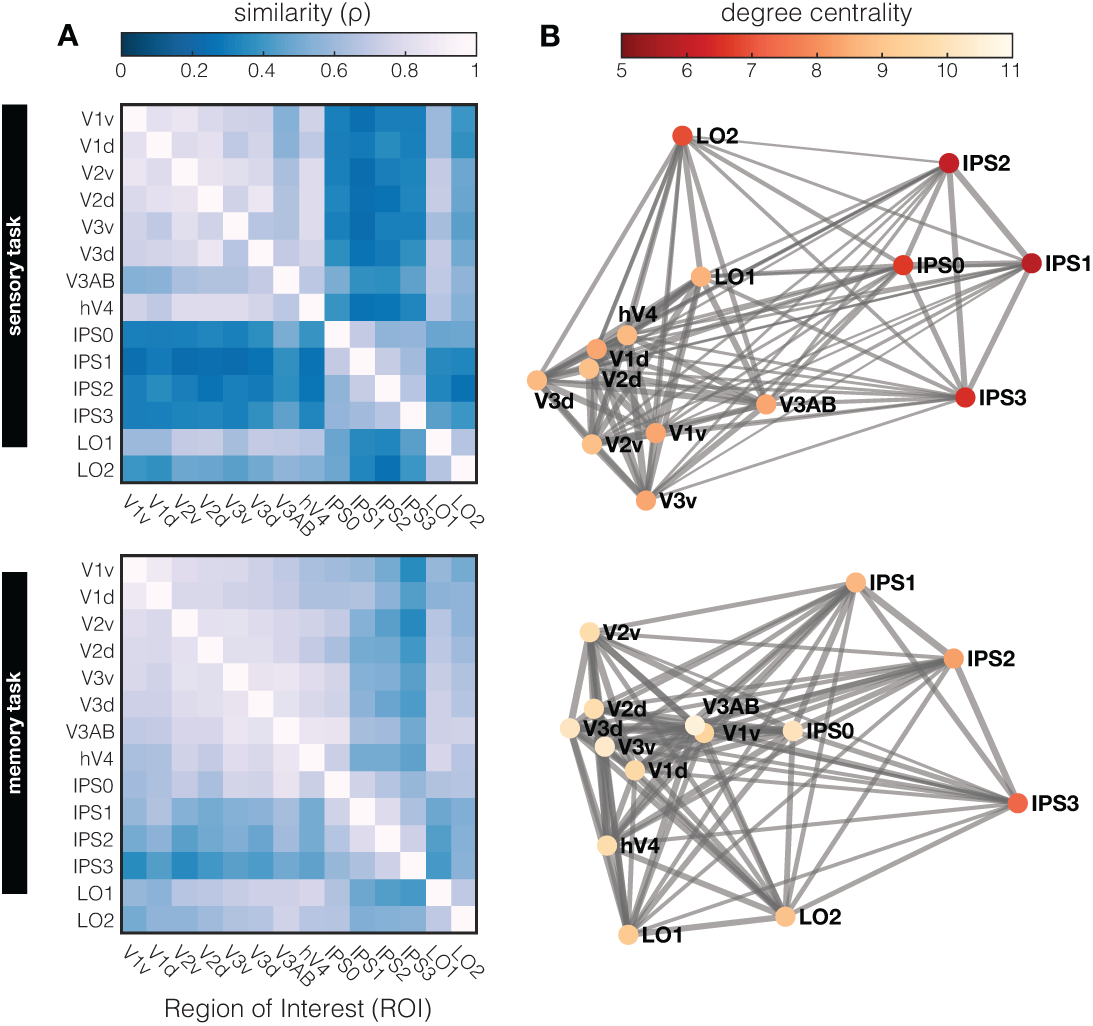
Second level RSA. **(A)** To compare how orientation is represented across different regions of visual cortex, RSM’s from fine-grained individual ROI’s (Supplementary Figure 1) were correlated in a 2^nd^ level similarity analysis. For the sensory task (top panel), representational similarity is high among early visual areas; high among the various IPS regions; and high among LO regions. Similarity between these three clusters is relatively low. For the memory task (bottom panel) there is a slight shift in similarity compared to the sensory task, with V1 becoming less similar, and IPS0 becoming more similar, to areas V2–V4. Furthermore, the distinction between areas is generally less pronounced. **(B)** Representational similarity can also be used as an indicator of connectivity between ROI’s based on shared representational geometry: When the geometry is similar, the “connection” is stronger (indicated here by the width of the grey lines connecting different ROI’s). The sum of the strength of these connections in a given ROI (i.e., degree centrality) indicates to which extent a local representational geometry resembles that of other ROI’s. Degree centrality is highest in early visual cortex and lowest in IPS regions, indicating a higher conservation of geometry across early visual cortical regions.

During the sensory task, there is notable shared representational similarity amongst early visual areas (V1–V4) and amongst areas in the intraparietal sulcus (IPS0–3), but low similarity between the two (Figure 5A, top). During the memory task, orientation geometries across various ROI’s show a somewhat different inter-areal organization (Figure 5A, bottom). First, the overall similarity between ROI’s is more pronounced during the memory task, with higher overall correlations between ROI’s compared the sensory perception task (r = 0.553 + 0.132 s.d., versus r = 0.32 + 0.082 s.d., respectively, with p < 0.001). This implies that there are fewer transformations of orientation geometry along the visual hierarchy during working memory compared to perception. Second, the cluster of early visual ROI’s with high representational similarity that was observed during perception (i.e., V1–V4), is shifted “upwards” along the visual hierarchy during memory – with V1 becoming less similar, and IPS becoming more similar to rest of EVC.

Finally, we probe the underlying “representational connectivity” structure in individual participants^46^. Unlike traditional functional connectivity analysis, the representational connectivity approach does not target covarying activation per se, but rather assumes connections on the basis of shared representational geometry. Visualizing these “connections” in a graph (Figure 5B) highlights the dense clustering of early visual cortex, weaker connections to LO, and weakest connections to IPS, both during the sensory and memory tasks. Based on this graph, we can compute the degree centrality of each ROI (or “node” in this graph) as the sum of connection strengths to other ROI’s. A high degree centrality denotes high representational similarity to many –or an especially strong representational similarity with some– other brain regions. Highest degree centrality is observed in early visual cortex, suggesting substantial overlap in representational geometry across these early regions. More downstream visual areas (IPS and LO) show the lowest degree centrality, implying a gradual transformation of representational geometry along the visual pathway that results in geometries not present at earlier processing stages. This analysis also shows how *functional* measurements of sensory driven responses to oriented gratings, and even responses during working memory when no stimulus is on the screen, are transformed along functionally defined ROI’s in a manner that is commensurate with the known *anatomical* structure of the visual hierarchy. This further highlights the power of an approach like RSA, whereby functionally measured response patterns are transformed into an abstracted representational space.

## Discussion

Here we compare how a simple visual stimulus is represented when it is either perceived or temporarily held in working memory, and show fundamental differences in the representational geometry of visual perception and visual working memory throughout human retinotopic cortex. By looking at the similarity of response patterns evoked by grating stimuli of different orientations, combined with a novel modeling approach, we are able to demonstrate relatively veridical representations during perception, and more categorical representations during working memory. We also find that the extent to which working memory representations are categorical increases along the visual hierarchy from posterior-to-anterior visual areas – modeling results that can be readily verified by simply looking at the geometries in the data. Importantly, our models make distinct predictions about veridical and categorical representations from a *single* input function based on the statistics of the natural world, which can also be implemented by measuring human behavior with a simple psychophysical task. This makes our modeling approach a potentially powerful tool to apply in other research contexts as well. With clear differences in representational geometry, it seems unlikely that working memory representations are merely noisier versions of perceptual representations, and our data imply a systematic compression of the coding scheme in parts of orientation space as a basis for categorization in working memory. Finally, by looking at inter-area representational similarity we recover known anatomical cortical structure, and observe a high degree of similarity for areas within early visual cortex (EVC), intraparietal sulcus (IPS), and lateral occipital cortex (LO) – but relatively low similarity between these respective regions.

Previous work from our group has claimed that visual working memories are represented in a “sensory-like” manner in early visual cortex^31^. This conclusion was drawn from the ability to cross generalize from sensory evoked responses, to responses recorded during the delay of a working memory task (using multivariate decoding techniques). However, there are multiple clues that VWM representations may be abstracted away from sensory evoked responses^30–32,63–66^, including the considerable differences between perceptual and working memory geometries unveiled in the present work. From a conceptual point of view there may be good reasons to keep formats distinct, as having identical representations for visual inputs and visual memories might make it difficult to distinguish external reality from internally generated thought^6,67,68^. Moreover, some sort of transformation of the information held in mind is often necessary to adequately support behavioral goals and motor output^69,70^. How can we reconcile the apparent contradiction between successful sensory-to-memory cross-generalization on the one hand, and the mounting evidence favoring abstraction during VWM on the other?

To understand how perception and working memory can evoke overlapping response patterns (i.e., have an overlapping *coding scheme*) while also differing in their representational format (i.e., the *representational geometry*), we will examine the relationship between multivariate decoding and RSA more closely. First, note that within any kind of task, a high degree of similarity along the diagonal of a cross-validated RSM is a prerequisite for successful multivariate decoding, and vice versa. After all, if a particular stimulus would evoke uncorrelated response patterns every time it is presented (i.e., no similarity along the diagonal), a decoder would not be able to predict such a stimulus from the disparate response patterns that make up its training set (i.e., no decoding). A lower off-diagonal similarity is a second prerequisite, as otherwise the patterns evoked by different stimuli are indistinguishable. Thus, in areas of the brain that care about a certain kind of stimulus, you can expect both a clear diagonal component in the RSM, as well as successful within-task decoding. Things are a bit more nuanced for continuously varying stimuli such as orientation. To illustrate: Two identical orientations might evoke similar patterns of responses, give or take some noise, but so will two orientations that are adjacent in orientation space. While two orientations that are further apart are likely to evoke dissimilar responses. Based on the gradual transition in physical similarity between continuously varying stimuli, one would predict an RSM pattern where similarity gradually drops off at larger distances from the diagonal. In this case, off-diagonal similarity can have meaning, albeit with diminishing returns as representational similarity decreases with increasing stimulus distance.

For our data, this means that the clear diagonal component in the RSM’s during both perception and working memory (see EVC ROI’s in Figure 1C) are indicative of the ability to decode orientation *within* each of these two tasks. However, such apparent overlap in the represenational geometry around the diagonals does not speak to the overall geometry, nor does it speak to the ability to decode *between* the sensory and memory tasks (via cross generalization). With respect to the overall geometry, we know that even relatively subtle transformations (e.g., shifts or warping) of an otherwise fairly stable underlying coding scheme can lead to dynamics in the low-dimensional geometry^39,71^. Vice versa, the representational geometry can remain stable in the presence of dynamics in the coding scheme^39–41^. With respect to cross-generalization, this means that a clear diagonal component in both perception and working memory RSM’s could in theory stem from totally different non-generalizable response patterns in one task compared to the other, as long as the pairwise distances between response pattern are comparable between tasks. We show that in early visual ROI’s the underlying coding schemes during the sensory and memory tasks are sufficiently similar to yield a clear diagonal component in an “across-task RSA” (Figure 4). Importantly, we validate our across-task RSA approach against a common implementation of multivariate decoding for continuous stimulus spaces (the so-called inverted encoding model^31,62^). More interestingly, the across-task RSM also provides some insight into how the coding scheme may be warped during working memory compared to perception in V1 – we observe biases away from cardinal orientations during working memory, with many remembered orientations resulting in response patterns that are similar to those of directly perceived oblique orientations. To sum up, working memory representations in EVC can be “sensory-like”, in that there is considerable overlap in the response patterns during perception and working memory. At the same time, a systematic warping of the geometry for orientation during the memory task, relative to the sensory task, may result in a more categorical geometry during VWM with high similarity around obliques.

In addition to using cross-generalization from sensory-to-memory responses to conclude that early visual areas store “sensory-like” working memory representations, our previous work drew upon the *lack* of such cross-generalization (in the presence of high within-task decoding performance) to conclude that IPS stores working memories in a format “transformed away” from sensory-like responses^31,72^. However, our current analyses reveal how the representational geometry during working memory is predominantly categorical *throughout* much of the visual hierarchy: A significant benefit of the categorical over the veridical model can be seen in V3– IPS0 when using a theoretical input function (Figure 2A, 2D bottom panel), and when using the psychophysical input function (Figure 3A, 3C bottom panel). The reason that cross-generalization from the sensory to the memory task fails in IPS may therefore not be due to a transformation in the representational geometry from earlier-to-later visual areas during working memory.

Instead, IPS might simply process information quite differently during perception than during working memory. Recent recordings from non-human primates reveal that the receptive field of an IPS neuron (in lateral intraparietal “LIP” cortex) that was demarcated by showing the animal visual stimuli on a screen does not necessarily overlap with the receptive field of that same neuron when demarcated by measuring responses during a delayed-match-to-sample task^73^. This implies a distinct mechanism for representing sensory inputs and working memory contents at the level of single neurons, which plausibly scales up to the level of population recordings as obtained with fMRI. A second reason why cross-generalization from perception-to-memory might be lacking in IPS is because the sensory input is represented rather weakly in IPS in our sensory task. While the full-field grating stimulus was attended, the feature of interest to our analyses (orientation) was not directly relevant to the participants’ task (detect contrast changes). This means that the sensory task signal-to-noise may have been insufficient for cross decoding. Alternatively, feature-based attention might change or improve stimulus representations in IPS, as it was shown to do in EVC^78^. Given the central role of IPS in attention^74–77^, it would be interesting to examine how attending different features of the same stimulus might impact stimulus representations, and whether this could explain the relatively noisy RSM’s we observed in IPS during our sensory task (Figure 1C, top).

Of course, the attentional state in different tasks matters more generally in terms of the conclusions we can draw. It’s been shown that a decoder trained on patterns of responses to perceived but unattended orientation stimuli (RSVP task at fixation) generalized to response patterns during the working memory delay^1^. By using *unattended* gratings, the authors could conclude that orientation-selective responses for remembered gratings depend on the same orientation-selective subpopulations driven during a bottom-up sensory response. In our work, we wanted to additionally control the deployment of spatial attention (presumably much narrower for a task at fixation) by having participants *attend the contrast* of a perceived grating. We already know that cross-generalized (sensory-to-memory) and within-task decoding (memory-to-memory) work well under these conditions^31^, so this choice meant we could be confident that possible geometrical differences would not be driven by differences in response patterns (or SNR) between the sensory and memory tasks. For future work it would be interesting to compare the sensory geometry resulting from our contrast attention task to a situation where the grating is not attended (task at fixation), or when the orientation of the grating is attended. We would speculate that in the presence of orientation attention, the geometry of a perceived grating would become even more veridical, as attention can be anchored to relevant attributes of the physical stimulus, spacing orientation representations more regularly throughout the 180° feature space.

A big question in the field of VWM concerns the role of primary visual area V1 during memory maintenance. On the one hand, sensory recruitment theory posits that involvement of area V1 is critical to maintaining highly detailed visual representations, as this is the only site thought to have the resolution to do so^21,79^. In support of this theory, many fMRI studies have shown that VWM contents can be decoded from V1^1,2^, and correlations between behavioral and decoding performance imply a functional role for V1^72,80^. On the other hand, outside of the fMRI literature there is less evidence to support a neural correlate of VWM in area V1, with a general failure to find sustained firing in EVC^81,82,83^ (but see also^84,85^), which has led some people to conclude that V1 decoding could be epiphenomenal^67,68^. Might our findings speak to this discrepancy between fMRI and single cell recording? Receptive fields in V1 are small, so if there is a representational shift from “veridical” during perception, to more “categorical” during VWM (presumably under the influence of top-down feedback), then working memory contents may be coded by a (subtly) different subset of neurons than those that respond to perceptual input. The reasoning that the same neurons may not code for the same stimulus under different task conditions holds true on several levels. For example, multi-unit activity associated with working memory maintenance was restricted to deep and superficial layers in V1, while such activity during perception also included the input layer 4^86^. Thus, even small shifts in the neural code (from one layer to the next, or from one orientation column to the next) may decrease the chance of finding sustained spiking when a one-to-one mapping between perception and working memory is assumed. Only looking at population wide neural coding, as we do here, can uncover working memory contents that has undergone a shift in representational geometry relative to perception.

Relying on population responses from fMRI BOLD does have one obvious caveat, which is the slow temporal profile of the signal. However, stimulus-evoked BOLD from the memory target can likely not explain the geometry during the working memory delay. Ample work has shown that even with identical sensory inputs at encoding, such BOLD does not carry stimulus information into the working memory delay when active maintenance of a stimulus feature is no longer required, with decoding performance for task-irrelevant features quickly dropping to chance^4,8,44,45^. For example, if one of two consecutive stimuli is cued for report immediately after stimulus presentation, the cued target cannot be decoded in the delay that follows^1^. When shown a stimulus with two independent features (i.e., orientation and color), only the remembered feature can be decoded during the delay^2^. Etc. Finally, clear geometrical differences between the sensory and memory RSM’s would not be expected if both tasks reflected the same response.

A well-known strength of RSA is that it projects response patterns associated with different task conditions (in our case, task conditions are 180 levels of orientation) into an abstracted space where representations can be compared between imaging modalities, species, models, or with behavior^46^. Thus, RSA is a powerful tool to sidestep the correspondency problem, allowing us to compare the output of systems that differ greatly. For example, one can construct an RSM from behavioral responses and correlate it with an RSM constructed from neural responses of specific brain regions^6,46,47^. However, visual perception involves a cascade of processes of increasing complexity, from simple feature-detectors in primary sensory cortex, to more invariant and category-based representations in ventral visual cortex. Behavior is the result of the entire brain working in concert to produce one output^87^, which means that even for a very simple stimulus such as orientation, the representational geometry can differ between areas and tasks. Using a behavior-derived RSM as a model could therefore miss a lot of variability in representational geometry across cortical areas, or produce misleading conclusions about a given area representing orientation while ignoring other areas that match behavioral output less. This is why using behavioral measurements (here: psychophysical input function) in a hypothesis-driven manner as the basis for multiple models (here: veridical and categorical), may allow for deeper insight into which specific areas of the brain are involved in multiple underlying components of a single behavior.

Our models are indeed able to quantify notable differences in representational geometry between orientation perception and working memory tasks, as well as differences between various retinotopic areas within a single task. However, there remain patterns in the data that neither of our current models are designed to capture. Most notably, data recorded during the sensory task from area V3AB (and arguably also V4, IPS, and LO) show a representational similarity pattern implying that the obliques (45° and 135°) are quite similarly to one another (Figure 1C, top row). This pattern hints at another possible form of (object-level) categorization, where all tilted lines, irrespective of their tilt direction, are represented in a similar fashion. This parallels evidence showing that people use the same verbal labeling (“diagonal”) for both obliques^88^. Alternatively, this pattern in V3AB could reflect something about the task, because tilt direction (45° or 135°) matters in the memory task due to the orientation recall requirement after the delay. During the sensory task, only contrast is attended. Precise top-down orientation signals during memory may override the tilt-agnostic tendency that certain areas (such as V3AB) might otherwise display.

Another example where we see the data diverge from our models, is in the non-uniformity along the diagonal of our RSM’s. Specifically, for the sensory task there appears to be lower similarity for horizontal (90°) compared to vertical (0°) orientations in many ROI’s, and similarity also appears relatively higher for obliques (45° and 135°) in some ROI (Supplementary Figure 6A). This non-uniformity implies that already at the level of sensory-driven responses all orientations are not represented equally, which is a well-established finding^59,60,89,90^. The higher overall similarity for oblique compared to cardinal orientations is exacerbated in the memory task (Supplementary Figure 6B). This may be due to a compression of orientation space around oblique orientations, leading to highly similar response patterns in this part of orientation space, while the pattern around cardinals is more distinct. This idea is supported by results from multidimensional scaling (Supplementary Figure 6C), which takes into account the entire geometrical pattern, showing stronger clustering around obliques compared to cardinals during the memory task.

Our categorical model was designed only to capture an extreme version of inhomogeneities in orientation space between cardinals and obliques. However, these inhomogeneities appear to some extent during the sensory task as well, and differ across the visual hierarchy. To directly test the extent to which these inhomogeneities appear across tasks and ROI, we designed an alternative model that uses a single parameter to modulate the magnitude of inhomogeneities (Supplementary Figure 7A). Specifically, by varying the exponent of the input function we create a family of model RSM’s that range from having higher similarity around cardinals (as also our veridical model), to diagonal, to higher similarity around obliques (including our categorical model). In fact, this approach closely approximates a previously proposed model^57^ that was not based on natural input statistics. When we fit this model to our data, we again show a gradual transition toward more categorical representations (with high similarity around obliques) along the visual hierarchy during memory, but not perception (Supplementary Figure 7B). Of course, we would construct even more complex models, with (many) more parameters, to eventually capture every specific geometrical pattern in every ROI. But this was never our goal. Instead, the main point of our models is to have a simple yet well-motivated quantification of the stark differences in geometry between the sensory and memory task, and the changes in the geometry along the visual hierarchy (in the memory task), that are readily observable by eye (Figure 1C). We show that these differences can indeed be quantified, and hold independent of the exact fitting approach (Supplementary Figure 4), binning approach (Supplementary Figure 2), and modeling approach (Supplementary Figure 7).

Given the diversity in RSM patterns described above, how might we compare the geometry within a given task, while remaining agnostic to the precise patterns in different regions of interest? To evaluate inter-area representational geometry differences in a more hypothesis-free manner, we used a “representational connectivity” analysis, which subsumes all possible patterns by simply quantifying degree of overlap. This allows us to compare how the visual system orchestrates representations across large swaths of cortex during both perception and working memory. One observation that emerges from this approach is that during VWM we see a shift in the interplay between areas, as compared to during perception. Specifically, during working memory the geometry in V1 becomes more differentiable from the geometry in the rest of EVC, and IPS0 becomes more differentiable from the rest of IPS (but more similar to EVC). In other words, we see that the inter-areal structuring of representational geometry differs between perception and memory. Another observation is that compared to perception, geometries during VWM show higher similarity across visual cortical areas. Such homogeneity might be expected if a unitary categorical top-down signal dominates feedback signals to multiple earlier areas. After all, in the absence of visual input, working memory information in V1 must be coming from within the brain itself, either through feedback connections or local recurrent processing.

Using representational similarity, in combination with the novel modeling approach outlined here, can be a powerful tool for studying representational formats during perception and working memory. Once the input statistics are known, either by deriving them from the environment or behavior, these models can be applied to any feature. Thus, in addition to tapping into a possible categorization for spatial features in EVC^32^, our approach has potential for surface-based features such as color or contrast, more complex visual objects such as shapes or faces, as well as stimuli in other sensory modalities. Many of the well-known advantages from RSA approaches also apply here, such as the potential to fit models across the entire brain in a manner that is not restricted to early sensory areas for which the receptive field mapping is known. Furthermore, the approach can be used with different measurement techniques that have a higher temporal resolution, allowing additional queries about the temporal progression of representational geometry as stimuli are encoded, remembered, and recalled.

## Methods

### Stimuli and procedures

We used a publicly available dataset, originally part of a study by Rademaker et al. (2019). This dataset contains both a visual perception task (the “sensory task”) and a visual working memory task (the “memory task”) performed in the scanner, and presented in the paper as Experiment 1 (6 participants with mean age = 28.67, sd = 3.675, and 5 females). The dataset also contains an independent psychophysical experiment, presented in the original paper as Supplementary Figure 9 (21 participants with mean age = 20.12, sd = 2.01, and 14 females. For this psychophysical experiment, only 17 participants were included in the analysis (3 dropped out, and 1 performed at chance level). All participants who contributed to the dataset were neurologically healthy, had normal or corrected-to-normal vision, received monetary reimbursement, and provided their written informed consent. Data were collected at the University of California San Diego.

In the scanner, both the sensory and memory tasks used full-contrast donut-shaped grating stimuli (1.5° and 7° inner and outer radius, respectively) with smoothed edges, a spatial frequency of 2 cycles per degree, random phase, a pseudo-randomized orientation. Stimuli were presented against a uniform grey background, and participants fixated a 0.4° central dot throughout. In the sensory task, donut-shaped gratings were presented in 9 second trials, contrast reversing at 5 Hz. Such donut trials were interleaved with trials showing a circular grating (1.5° radius), and fixation periods (10% of total). Grating contrast was briefly (200ms) and probabilistically reduced to 80% Michelson twice every 9s, and participants’ task was to report such contrast changes. Participants completed a total of 300-320 sensory task trials across 3 separate scanning sessions. The sensory task was also used to localize visually responsive voxels (via a donut > circle contrast), and in our current analysis we use this contrast as a mask for all EVC ROI’s (but not IPS and LO, where all retinotopically defined voxels are included). In the working memory task, a target grating was shown for 500ms, and recalled 13 seconds later by rotating a white line (spanning 7°) for 3 seconds to match the remembered orientation. Between trials, there could be 3, 5, or 8 second fixation intervals. During the delay of two-thirds of memory task trials, a distractor of 50% Michelson contrast could be presented for 11 seconds during the middle portion of the delay. Distractors could be a grating (1/3^rd^ of trials) or filtered noise (1/3^rd^ of trials), contrast reversing at 4Hz. By ensuring uniform orientations of grating distractors with respect to memory targets, we are able to look at the representations of remembered and distractors orientations independently. Importantly, due to the negligible differences between trials with or without distractors, both in terms of behavior as well as decoding, we collapse the data across all working memory trials for our main analyses. Each participant completed 324 total working memory trials over the course of 3 different scanning sessions.

The independent psychophysical experiment was completed outside of the scanner, and stimuli consisted of gratings presented at 20% Michelson contrast against a uniform grey background. Gratings had a 2° radius, spatial frequency of 2 cycles per degree, random phase, and pseudo-randomized orientation. Each trial started with a 200ms target orientation that was remembered over a 3s delay, and recalled by rotating a dial to match the remembered orientation in an unspeeded manner. Intervals between trials were 800-1000ms. Grating distractors were presented for 200ms during the middle portion of the delay on 90% of trials. As in the scanner, targets and distractors were uncorrelated across trials, allowing for independent analysis of responses to the target. Any biases resulting from distractor presentation were small (Supplementary Figure 9 of ref^30^) and did not exert much influence on responses. Each participant completed 1620 trials over the course of several testing sessions.

For more detailed information on scanning and task procedures, please reference Rademaker et al. (2019).

### Models

First, the function that constrains both the veridical and categorical models, and emulates the frequency distribution of orientations in the natural world, is described by

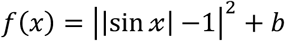

Where −π < *x* < π, and *b* is any non-zero baseline (due to z-scoring before fitting this function is scale-free). This function is loosely based on the function defined in ^38^. Another way to think about this theoretical “input statistics” function, is as the normalized amount of Fisher information at each orientation in orientation-space. We ensured that the results of our model fits were robust to the specific shape of the input function by not only using a theoretical input function based on the statistics of the natural world (Figure 2A), but by also using a psychophysical input function (generated from independent psychophysical measurements, Figure 3A) as the basis for our two models. Irrespective of the input function used (theoretical based on previous literature, or psychophysical based on independent data), model generation, as described next, is identical. Note, for the alternative model in Supplementary Figure 7, this function is also used, with the difference that the exponent (here set to 2) is a free parameter over range [0 inf).

For the veridical model, we assume a set of idealized tuning functions (Figure 2B, top) and use it to simulate neural responses to all possible stimulus orientations. From these simulated responses we calculate the similarity (rho) between all possible pairs of stimulus orientation (as shown in Figure 1B), resulting in a veridical model RSM (Figure 2B, bottom). Each tuning function in the veridical model is defined by a von Mises (circular analogue of a Gaussian distribution),

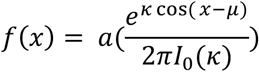

with a fixed concentration parameter *κ* = 6, a center defined by *μ*, and −π < *x* < π. *I*_+_(*k*) is the modified Bessel function of order 0. The amplitude *a* of each tuning function varies across orientation space as determined by the height of the input statistics function (i.e., from 0.1–1.1).

For the categorical model, we calculated the psychological distance between all possible pairs of orientation. For any pair of orientations, we sum over the approximated derivatives between these two points along the input statistics function as follows,

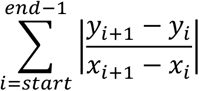

where *x* is an orientation in orientation space (and wraps around a circle), and *y* is the amount of normalized y-axis information for that orientation.

The veridical and categorical models described above were converted from radians to degree (spanning the entire orientation space from 1° to 180°, in steps of 1°) to match the dimensions of the data (also in degree). To evaluate if the representational structure in the data RSM’s (Figure 1C) is more or less similar to veridical or categorical model RSM’s (Figure 2B-C), both data and model RSM’s were normalized before fitting. The correlation between each model with the data RSM’s was done via semi-partial correlations, because the veridical and categorical models are not independent. Specifically, to evaluate the correlation between model A and the data RSM of a given ROI (*RSM_ROI_*), we first removed the variance explained by model B (*RSM^B^*) from model A (*RSM^A^*), to get its model residuals (*ε^B^*):

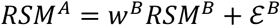

Where *w^B^* are the initial weights of model B, and the residuals *ε^B^* reflect any pattern in model A unaccounted for by model B. To illustrate, imagine the extreme case where model B explains none of the same variance as model A, then *w^B^* would be 0, and the residuals *ε^B^* would be equal to model A itself. Next, the residuals of model B are correlated to the data *RSM_ROI_*, and we apply a Fisher transformation (also known as inverse hyperbolic tangent, or *tanha^-1^* to normalize the bound correlation data (from –1 to 1), to unbound values (–Inf to Inf) suitable for statistical testing.

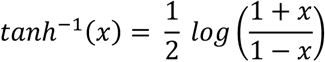

This transformed correlation gives the amounts of variance explained by model A independent of model B. To summarize, this procedure ensures that any resemblance (big or small) between the two models is accounted for first, after which we obtain the unique contribution made by model A.

To ensure that our results did not depend on the specific fitting approach described above, we also fit our two models directly to the data without taking into account the overlap between the models. The difference in model fits is still informative in this case (showing which model fits the data better), which was confirmed by statistical replication of the main findings. Second, we used general linear regression to fit both models simultaneously in the form of:

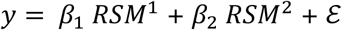

Where *β* reflects the beta weight for each of our two model RSM’s. Also using this procedure, the statistics of our effects replicated. For a figure showing both these approaches, see Supplementary Figure 4

### Across-task RSA fidelity

To directly relate cross-generalization from decoding (or more specifically, from the inverted encoding model, or “IEM”, as used in ref^30^) to our novel “across-task RSA” (Figure 4), we calculate a fidelity metric to quantify how much information there is about the remembered orientation based on the pattern responses to the perceived orientations. Because orientation is a continuous variable, we also take into account the representational similarity to orientations nearby the remembered one, and the expected drop in similarity at increasingly larger distances in orientation space. For each correlation profile (see Figure 4B–D) we calculate fidelity in a manner identical to how this has been calculated for IEM channel reconstructions (in ref^30^”, and shown here as the y-axis in Figure 4D). Specifically, we take the correlation value at each degree in orientation space (wrapped onto a 2π circle), and project this vector onto the remembered orientation (centered to zero degrees) via 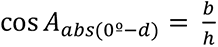, where *A* is the angle between the remembered orientation (at *0°*) and the degree in orientation space being evaluated (*d*), and *h* is the correlation value at *d* (i.e. the hypotenuse of a right triangle). This procedure was repeated for all 180 degrees in orientation space, and we then calculate the mean of all 180 projected vectors. This fidelity metric captures the amount of information at the remembered orientation, and removes additive offsets.

### Analyses of 2^nd^ level RSM and representational connectivity

To compare the representational geometry across all retinotopic ROI’s during perception and working memory, we employed two approaches first described by^41^: A 2^nd^ level RSA and a ‘representational connectivity’ analysis. Note that for these analyses we used the smallest possible ROI’s that were retinotopically defined, meaning we split early visual areas into their dorsal and ventral parts, and used the individual sub-areas of IPS and LO. For the 2^nd^ level RSA, we calculated the similarity of each across-subject RSM (as shown in Supplementary Figure 1) to every other across-subject RSM using spearman correlation (as RSM’s are monotonically, but perhaps not linearly related). This resulted in the 2^nd^ level RSM’s in Figure 5A, showing representational similarity between all retinotopic ROI’s during perception, and during working memory. For the ‘representational connectivity’ we similarly computed Spearman correlations between ROI’s but at a within-subject level as is the recommended procedure^41^. Across-subject averages are visualized in a graph (Figure 5B) where each node signifies a ROI, and each edge signifies the correlation coefficient to each other ROI. Thicker and shorter edges indicate higher similarity. We computed degree centrality for each node as the sum of all edges, depicted by the saturation of each node (less saturation indicating higher degree centrality). Thus, higher degree centrality indicates that a ROI shares its representational geometry either strongly with a few, or somewhat strongly with many, other ROI’s.

## Statistics

First, for the main analyses (Figures 2 and 3) we tested if model fits differed between task (sensory or memory), model (veridical or categorical), or ROI’s by running a three-way repeated measures ANOVA using R (version 4.1.1) and RStudio (version 1.4.1717). Significant three-way interactions (between task, ROI, and model) were followed up with two targeted two-way repeated-measures ANOVA’s – one for the sensory task, and one for the memory task (as described in the main text). Significant interactions arising from two-way ANOVA’s were further examined via post-hoc two-sided paired-sample t-tests within each task and ROI (uncorrected for multiple comparisons). For the main model fitting results using the theoretical and behavioral input functions (Figure 2D and 3C, respectively) these t-tests are reported in detail in Supplementary Table 1.

For the second order RSA (Figure 5) we tested if there was a significant difference between perception and working memory in terms of the overall correlations between ROIs (Figure 5A). First, we calculated the mean correlation between all ROI’s (excluding the diagonal) for a given subject within each task. During the sensory task the mean correlation between all ROI’s was r = 0.32 (+0.082 sd), and during the memory task it was r = 0.553 (+ 0.132 sd). Next, we performed a permutation test where we shuffled the assignment of each ROI-ROI correlation pair (i.e., each value in the 2^nd^ order RSM, excluding the diagonal) to either the sensory or memory task at random for each subject. On each permutation we then calculated the mean correlation between all ROI’s within the shuffled sensory and shuffled memory RSM’s, and took the difference between these means. A null distribution of differences was generated over 1000 permutations, and compared to the real difference between the mean correlations (i.e., 0.553 – 0.32 = 0.233), which was significantly greater than expected by chance (p < 0.001).

## Code accessibility

The data are public and can be accessed via the Open Science Framework (OSF) at https://osf.io/dkx6y which has an accompanying wiki. The code for the analyses in this paper can be found at https://osf.io/ej9db/.

## Acknowledgements

Data acquisition was financed by NEI R01-EY025872 and NIMH R01-MH087214 to John Serences – whose unwavering and continued support has shaped us as scientists. RLR and MDH are funded by the Max Planck Society, MDH is also funded by the German Federal Ministry of Education and Research (BMBF). We thank the IMP lab at Goethe University & Rademaker lab members at ESI for feedback on the manuscript. We also thank Xue-Xin Wei and Tim Brady for seeding the thoughts that changed our minds, culminating to the modeling work in this paper.

---

**Supplementary Figure 1:**
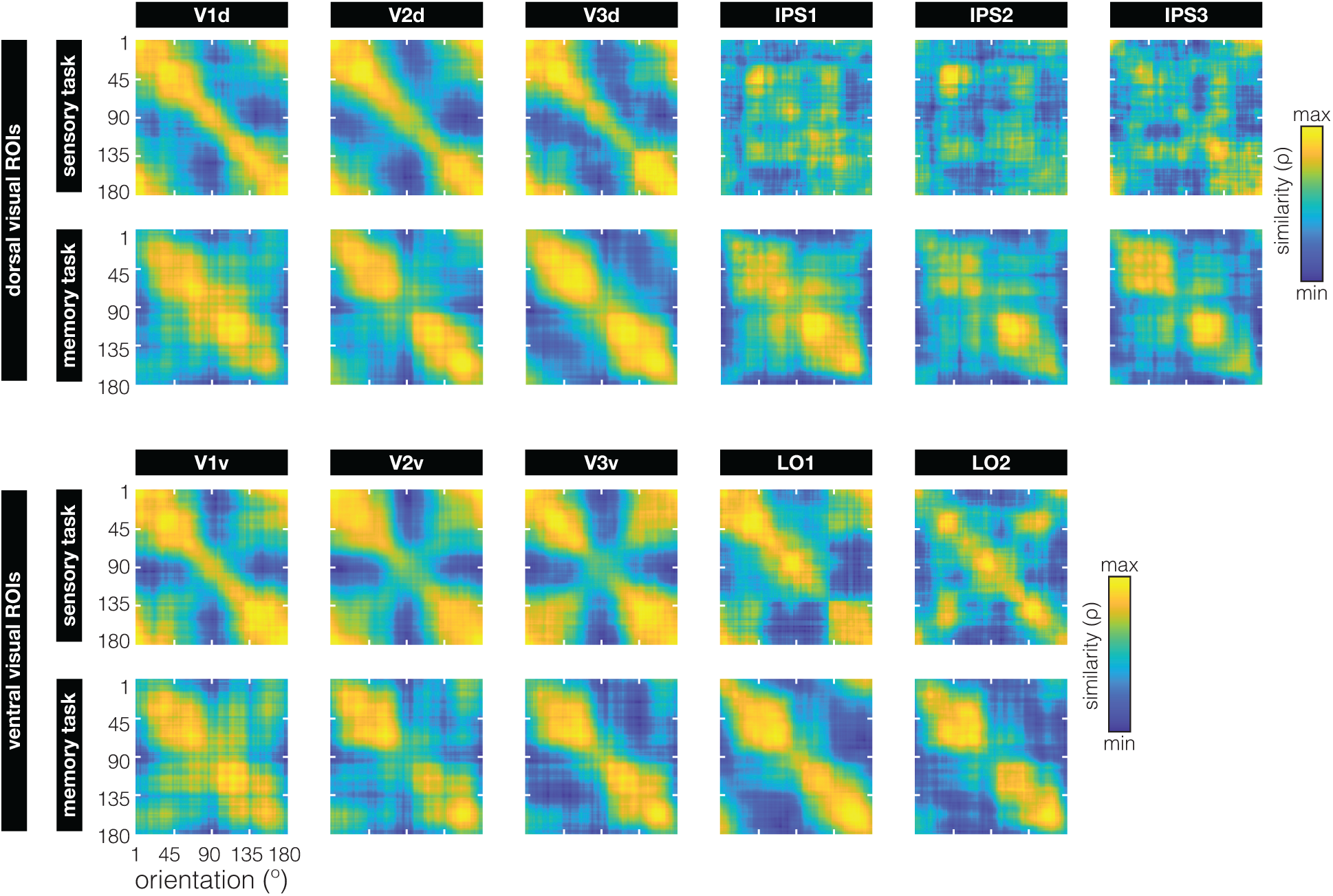
Orientation representational geometry (as indexed by RSM’s) during sensory perception and working memory for all retinotopically defined ROI’s (across all participants) that were not already shown in Figure 1C. Here, ROI’s are organized by whether they are located in the dorsal or ventral stream (top and bottom two rows, respectively). Early visual areas V1–V3 were split by their dorsal and ventral portions – used as input to the second-level RSA analysis (Figure 5 of the main text). Areas IPS1–3 (in the dorsal stream) and LO (in the ventral stream) were split based on their respective sub-portions – and similarly used as input to the second-level RSA analyses. All RSM’s are scaled to the range of correlations within each subplot to ease visual comparison of representational structure.

**Supplementary Figure 2:**
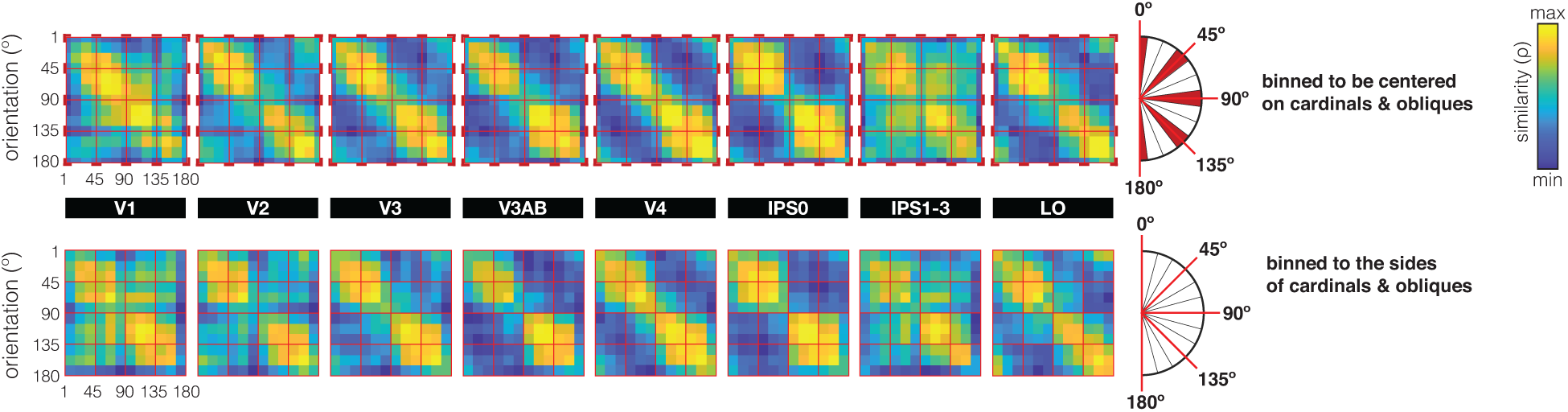
With 180 possible target orientations, and a finite number of trials, some form of smoothing or binning is necessary for RSA to yield reliable correlations. In our main analysis we smooth over a window of +10°, which could in theory impact the geometry around categorical boundaries (90° and 180°). In particular, it could induce some smearing of the categorical pattern observed in the memory task. To ensure that this pattern does not critically depend on the way trials are combined, here we show the data for the memory task binned (instead of smoothed) into 12 bins of 15°. On top, we show RSM’s with bins *centered on* the 2 cardinals and the 2 obliques (see inset), meaning that the parts of orientation space highlighted in dark-red are bins that include a cardinal or an oblique orientation. On the bottom, we show the same analysis but with the bins shifted, such that they respect cardinal and oblique boundaries, and bins *fall on either side*. We observe that similarity in bins that include a cardinal (top row) is relatively low, presumably due to the relatively large psychological distance between orientations on different sides of a cardinal orientation, resulting in lower correlations. Nevertheless, there is relatively low similarity around cardinals *also* when we respect the categorical boundary (bottom row), implying these categorical effects are not impacted much by the specific binning approach. Overall, binning or smoothing do not drastically change the geometry (though of course, the resolution of the RSM is much lower with binning).

**Supplementary Figure 3:**
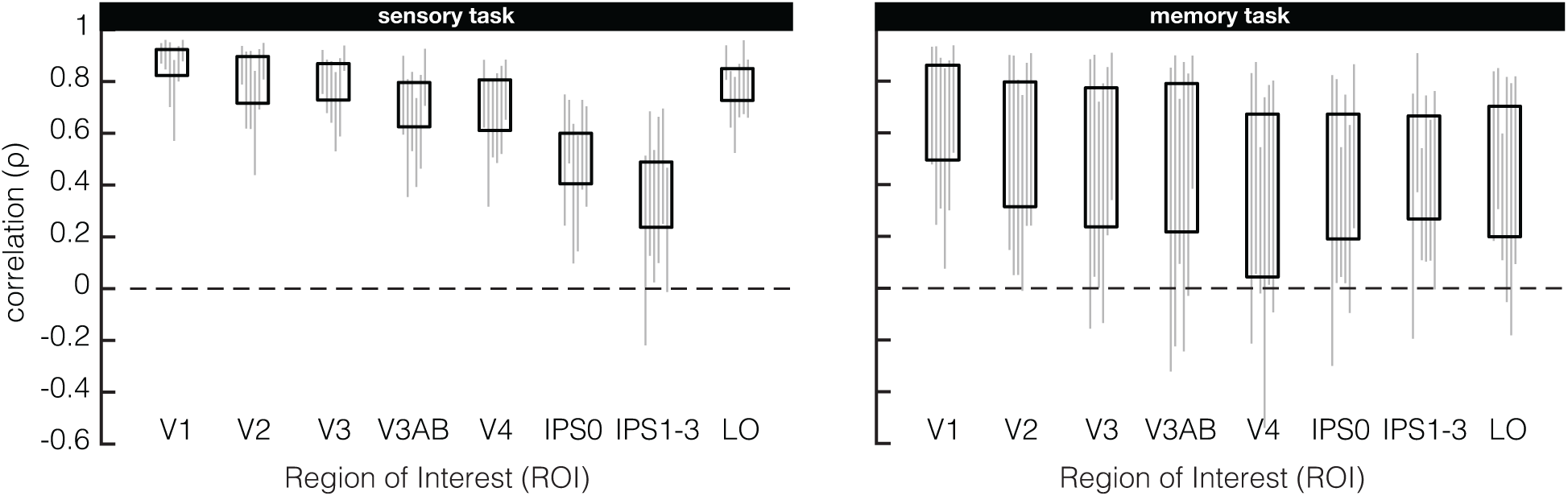
Exact ranges of correlations in the RSM’s from Figure 1C. To best show the representational structure for sensory and memory representations across ROI’s, and to ease comparison between them, the RSM’s in Figure 1C are scaled to the range (min-to-max) of correlations within each subplot. But the minimum and maximum correlations are not identical across subplots, therefore, correlation ranges across all participants (black rectangles) and individual participants (grey lines) are shown here for sensory (left) and memory (right) RSM’s.

**Supplementary Figure 4:**
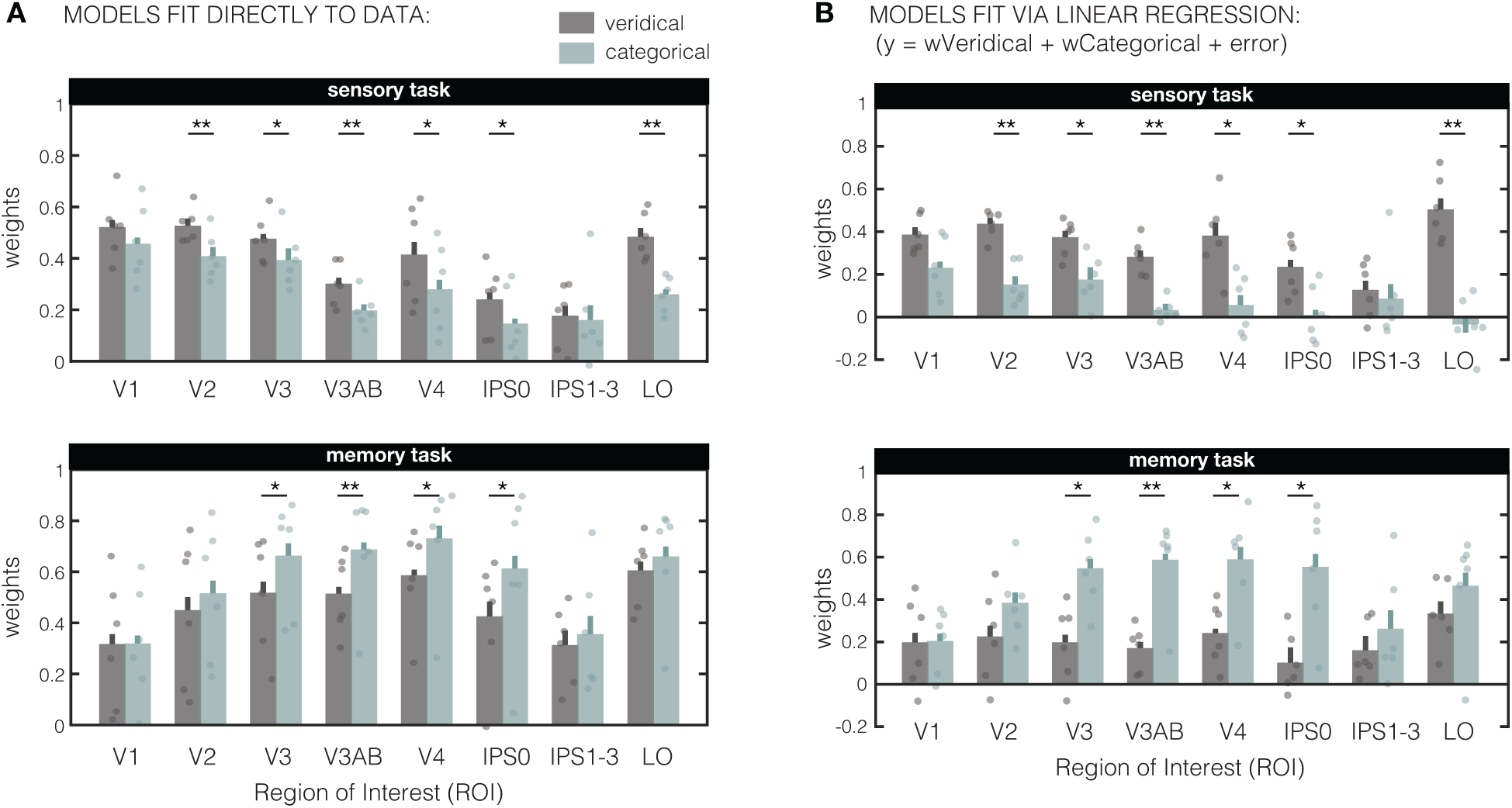
Two alternative fitting approaches. **(A)** Model weights when fitting the veridical and categorical models directly to the RSM’s (without first taking the residuals), and **(B)** model weights derived with a general linear regression (independent weights for each model). Irrespective of the fitting approach, the geometrical differences between our two tasks are captured by higher “veridical” weights in the sensory task, and more “categorical” weights in the memory task. For the “direct fitting” approach (in **A**) there is a significant 3-way interaction (model x ROI x task, F_(7,35)_ = 2.413; p = 0.0398), which we followed up by post-hoc ANOVA’s within the sensory and memory task separately. There is a main effect of model in both the sensory (F_(1,5)_ = 40.26; p = 0.001) and memory (F_(1,5)_ = 12.47; p = 0.017) tasks that is not the same in all ROI’s (as indexed by model x ROI interactions for sensory F_(7,35)_ = 3.262, p = 0.009 and memory F_(7,35)_ = 2.791, p = 0.024 tasks). Similarly, for the general linear regression approach (in **B**) there is also a significant 3-way interaction (model x ROI x task, F_(7,35)_ = 2.414; p = 0.0398), and main effects of model in both the sensory (F_(1,5)_ = 40.28, p = 0.001) and memory (F_(1,5)_ = 12.48, p = 0.017) tasks, and this difference between the models is not the same in all ROI’s (as indexed by model x ROI interactions for both sensory F_(7,35)_ = 3.26, p = 0.009, and memory F_(7,35)_ = 2.79, p = 0.024 tasks). Asterisks indicate the significance level of post-hoc two-sided paired-sample t-tests (*p < 0.05; **p < 0.01) comparing the two models in each ROI.

**Supplementary Figure 5:**
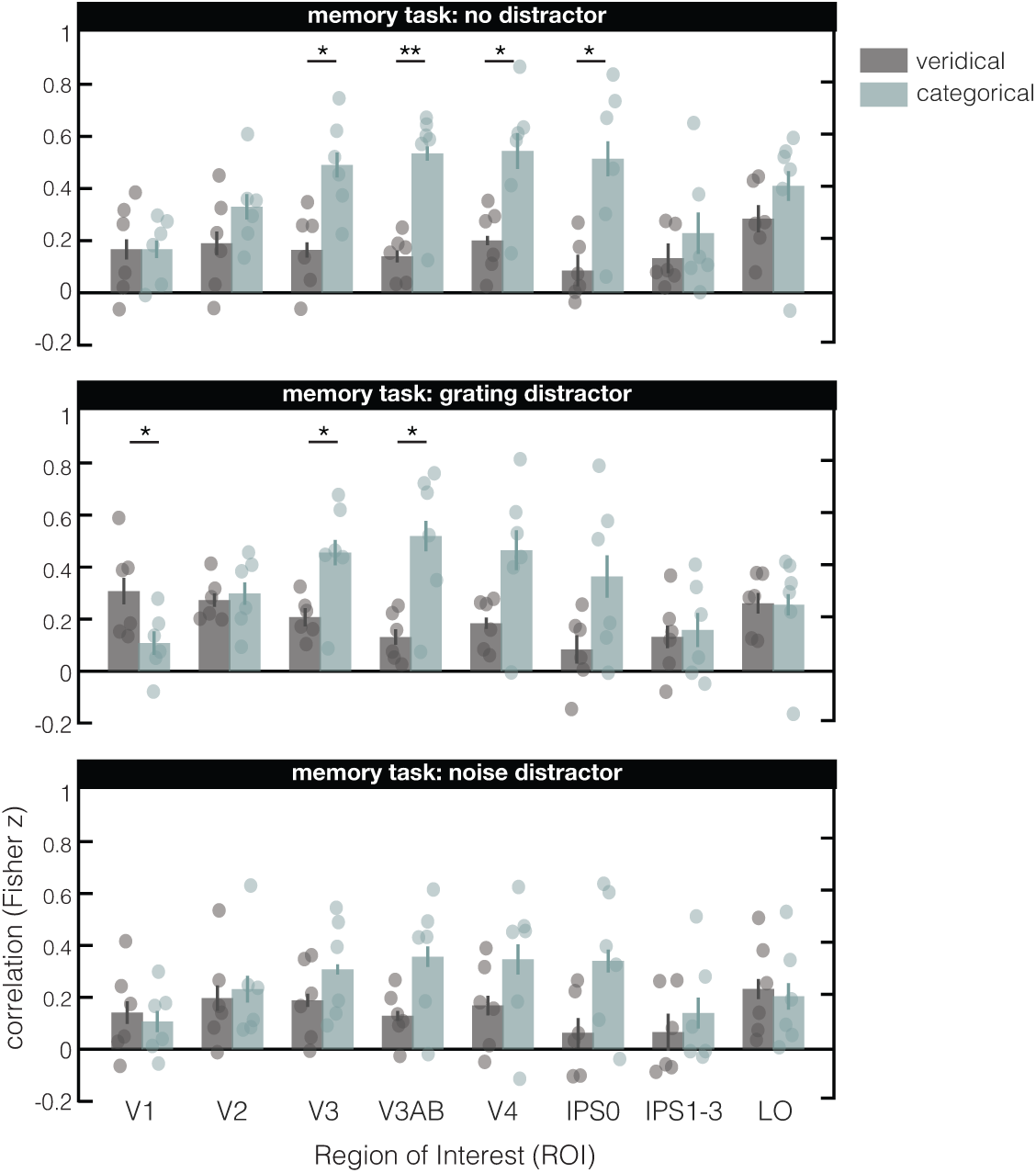
Model fits for the 3 different working memory distractor conditions. Overall, the results split by condition are qualitatively similar to the main results across all trails (Figure 2D, bottom panel). Two-way ANOVA’s comparing model and ROI showed that the categorical model did a better job at explaining the data in some of the ROI on trials without a distractor (model x ROI interaction: F_(7,35)_ = 2.914; p = 0.016; main effect model: F_(1,5)_ = 13.07; p = 0.015) and with a grating distractor (model x ROI interaction: F_(7,35)_ = 4.3; p = 0.0016; main effect model: F_(1,5)_ = 6.344; p = 0.053), indicating increasing differences between the veridical and categorical models along the visual hierarchy. While we see similar trends for the 108 trials with a noise distractor, these effects did not reach significance (model x ROI interaction: F_(7,35)_ = 1.812; p = 0.116; main effect model: F_(1,5)_ = 1.335; p = 0.3). Nevertheless, despite using only 1/3^rd^ of the data in each of these sub-plots, the pattern in the data is highly consistent. Asterisks indicate the significance level of post-hoc two-sided paired-sample t-tests (*p < 0.05; **p < 0.01) comparing the two models in each ROI.

**Supplementary Figure 6:**
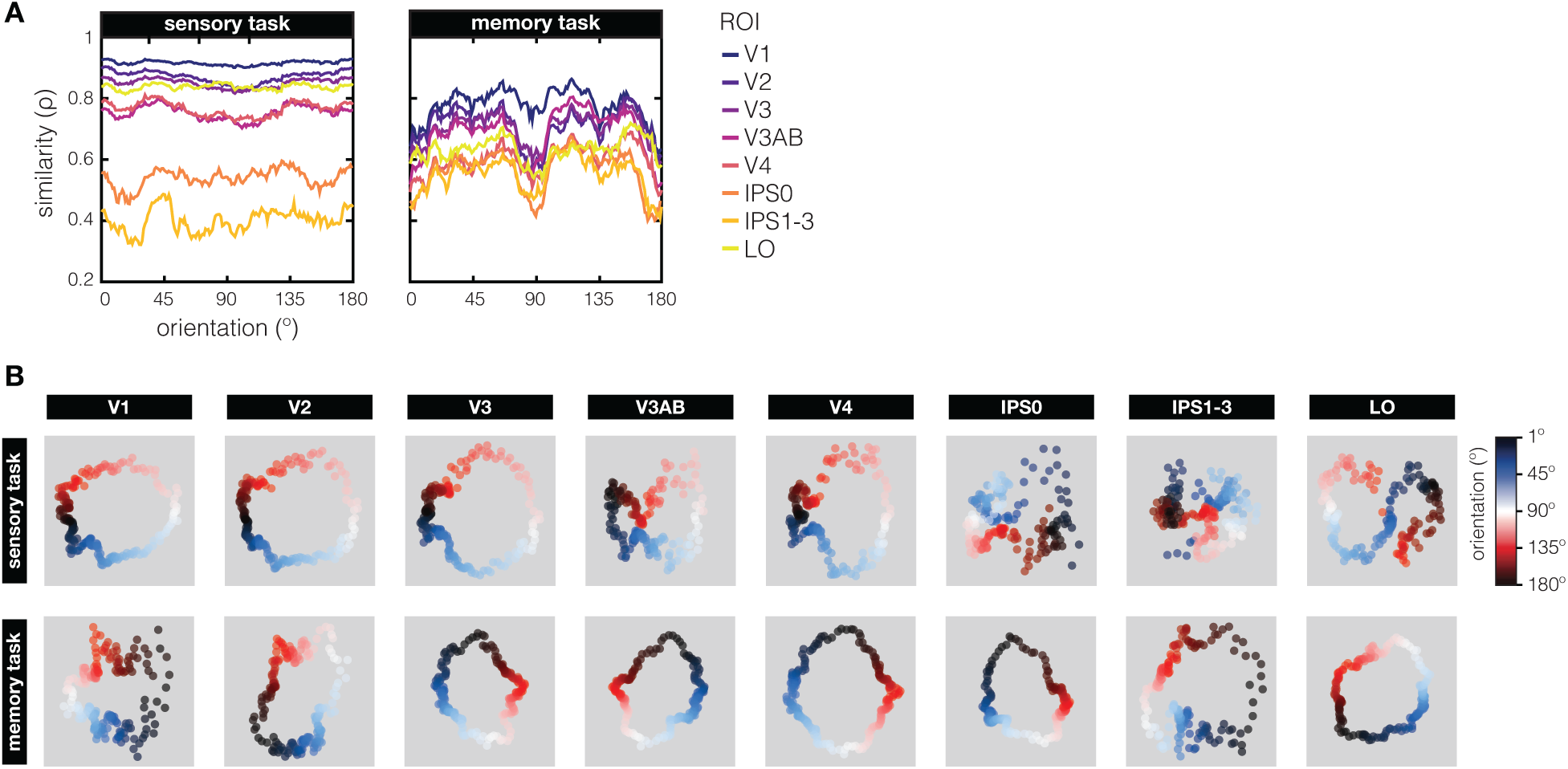
Orientation inhomogeneities of the representational geometry **(A)** To examine the inhomogeneity or representational similarity throughout orientation space, we plot the diagonals of the RSM’s from in Figure 1C. During the sensory task, we see that similarity tends to be relatively high around vertical orientations (0°/180°) compared to horizontal orientations (90°). For both tasks, oblique orientations are represented relatively more similar, and cardinals less similar. This “oblique” like effect is much exacerbated in the memory task compared to the sensory task. **(B)** We use multidimensional scaling (MDS) to projects high dimensional response patterns into 2 dimensions, in order to better visualize of how orientation space is represented. During the sensory task there is an orderly geometrical progression of orientation space, with the highest similarity between adjacent orientations (and some clustering around cardinal orientations, especially 180°) in early visual areas V1–V3. There’s also a “pinching” of orientation space around the obliques (45° and 135° become very similar) in more anterior visual areas V3AB–IPS and LO. During the memory task, the orientation space geometry remains circular in all ROI’s, with notable clustering of similarity around the obliques.

**Supplementary Figure 7:**
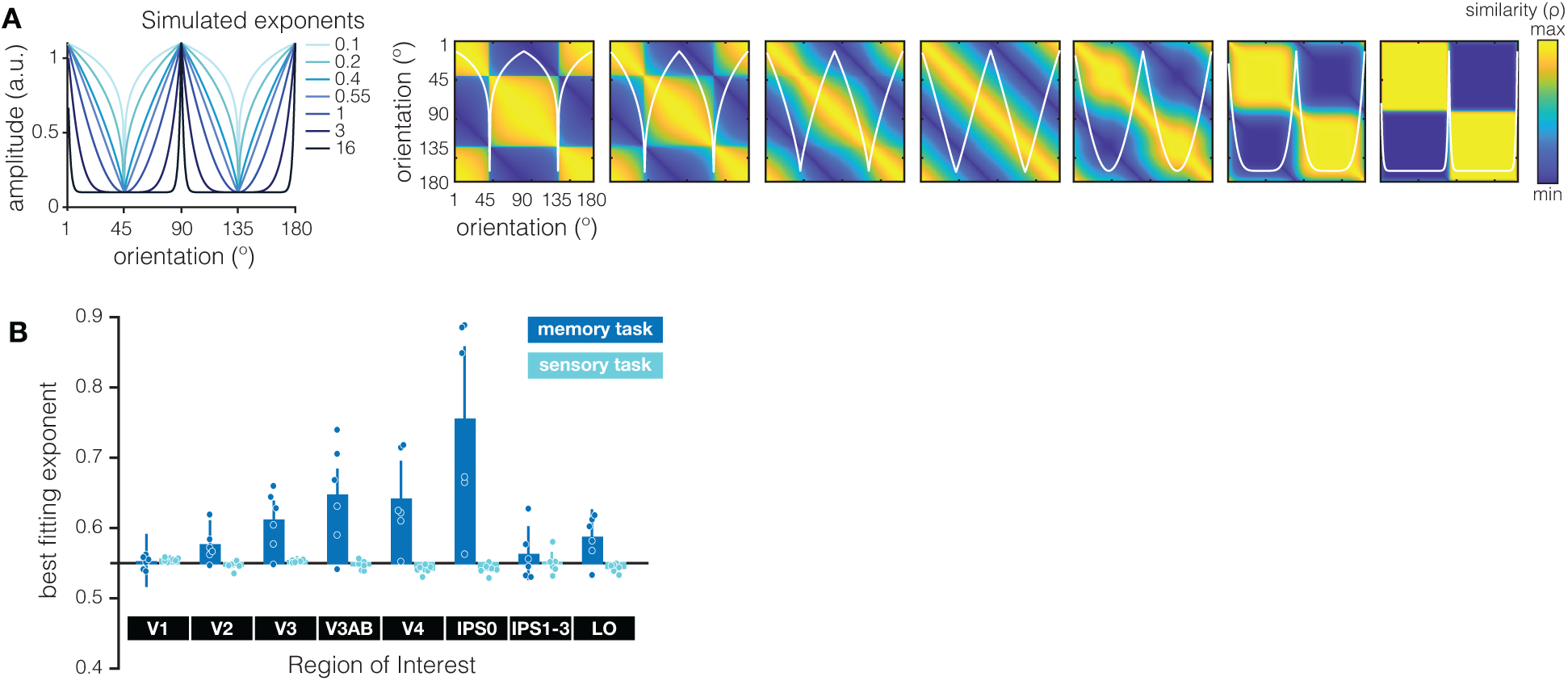
Alternative model based on psychological distance. **(A)** An alternative way to model the sensory and memory task RSM’s is to vary the degree of similarity that can be expected at cardinals or at obliques. By changing the exponent in the input statistics function *f*(*x*) = &|sin *x*| −1&^*exponent*^ + 0.1 to a free parameter, and using the psychological distance between every pair of orientations (as in the categorical model), we can create a family of input statistics functions (left panel) that modulate the shape of the model RSM’s (right panels) such that we can span any orientation anisotropy ranging from highest similarity around cardinals to highest similarity around obliques (similar to the modeling approach in ^57^). Each of the input functions in the left panel matches an RSM in the right panels (with the input function overlaid in white). At an exponent of 0.55 we approximate a uniform diagonal RSM, or a “physical” model of orientation space. Note that by modulating the shape of the input function in this manner, we can retrieve models that look very similar to our veridical model (e.g., exponent = 0.4), and a model identical to our categorical model (exponent = 2) in this parametric RSM space. **(B)** We plot the best fitting exponent for the input function for all ROI’s (x-axis) and separately for the memory (dark blue) and sensory (light blue) tasks, and show that those differ significantly (ROI x task interaction: F_(7,35)_ = 9.658; p < 0.001). For the sensory task the best fitting exponent stays close to 0.55 for all ROI’s, indicating that an RSM with a close-to uniform diagonal fits the data well. That said, the exponent does differ across ROI’s (main effect of roi, F_(7,35)_ = 3.307; p = 0.008), showing that a model with slightly higher similarity around obliques (exponent > 0.55) does better in for example V1, while a model with slightly higher similarity at cardinals (exponent between 0 and 0.55) does better at for example V4. For the memory task we see a gradual increase in the exponent along the visual hierarchy (up to and including IPS0), indicating that a model with increasingly stronger similarity around oblique orientations (i.e., increasingly stronger categorization) is better at explaining the memory geometry for more anterior ROI’s (main effect of roi, F_(7,35)_ = 9.213; p < 0.001). Best fitting exponents for individual subjects are shown as dots, and error bars indicate + 1 within-subject SEM.

**Supplementary Table 1:**
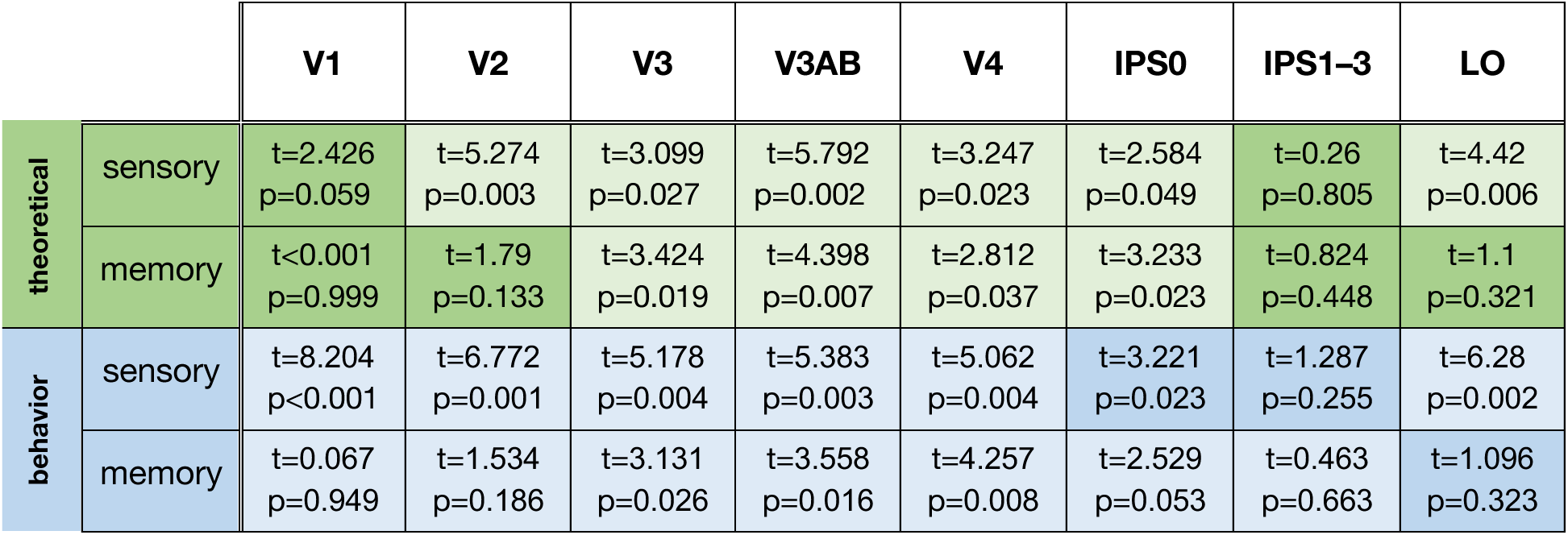
Post-hoc statistics for two-sided paired t-tests from the theoretical input function based on the statistics in the natural world (in green) and from the psychophysical input function based on independent behavioral measurements (in blue). All significant cells are colored in a lighter shade for the purpose of quick visualization.

|  |  | V1 | V2 | V3 | V3AB | V4 | IPS0 | IPS1-3 | LO |
| --- | --- | --- | --- | --- | --- | --- | --- | --- | --- |
| theoretical | sensory | t=2.426<br>p=0.059 | t=5.274<br>p=0.003 | t=3.099<br>p=0.027 | t=5.792<br>p=0.002 | t=3.247<br>p=0.023 | t=2.584<br>p=0.049 | t=0.26<br>p=0.805 | t=4.42<br>p=0.006 |
|  | memory | t<0.001<br>p=0.999 | t=1.79<br>p=0.133 | t=3.424<br>p=0.019 | t=4.398<br>p=0.007 | t=2.812<br>p=0.037 | t=3.233<br>p=0.023 | t=0.824<br>p=0.448 | t=1.1<br>p=0.321 |
| behavior | sensory | t=8.204<br>p<0.001 | t=6.772<br>p=0.001 | t=5.178<br>p=0.004 | t=5.383<br>p=0.003 | t=5.062<br>p=0.004 | t=3.221<br>p=0.023 | t=1.287<br>p=0.255 | t=6.28<br>p=0.002 |
|  | memory | t=0.067<br>p=0.949 | t=1.534<br>p=0.186 | t=3.131<br>p=0.026 | t=3.558<br>p=0.016 | t=4.257<br>p=0.008 | t=2.529<br>p=0.053 | t=0.463<br>p=0.663 | t=1.096<br>p=0.323 |

